# A Systematic Comparison of Single-Cell Perturbation Response Prediction Models

**DOI:** 10.1101/2024.12.23.630036

**Authors:** Lanxiang Li, Yue You, Yunlin Fu, Wenyu Liao, Xueying Fan, Shihong Lu, Ye Cao, Bo Li, Wenle Ren, Jiaming Kong, Shuangjia Zheng, Jizheng Chen, Xiaodong Liu, Luyi Tian

## Abstract

Predicting single-cell transcriptional responses to perturbations is central to dissecting gene regulation and accelerating therapeutic design, yet the field lacks a rigorous, task-spanning assessment of model behavior. We present a large-scale benchmark of 13 representative methods and baselines across 25 datasets spanning diverse perturbation modalities and species, including two primary immune-cell drug-response resources. We evaluated three core tasks—generalization to unseen single-gene perturbations, prediction of combinatorial interactions, and transfer across cell types—using 24 metrics covering expression-level accuracy, relative changes, differential expression recovery, and distributional similarity. Across tasks, performance depended strongly on perturbation effect size and evaluation perspective: expression-level agreement was highest for small-effect perturbations resembling controls, whereas delta- and DE-based metrics improved with larger effects, providing clearer signals. Models shared a conservative bias, with fine-tuned foundation models compressing variance and underestimating synergistic effects in combinations. PerturbNet showed superior recovery of DE signatures in Tasks 1 and 2, while no method consistently generalized across cell types in Task 3, where biological consistency dominated outcomes. This benchmark establishes current methodological limits, clarifies that different metrics probe distinct biological signals rather than redundant summaries of the same prediction problem, and provides a foundation for developing virtual-cell models that more faithfully capture heterogeneous perturbation responses.

**Teaser:** A large-scale benchmark reveals strengths and limits of models predicting single-cell perturbation responses.

## Introduction

Single-cell RNA sequencing (scRNA-seq) offers unprecedented resolution for exploring cellular heterogeneity (*1, 2*). Multiplexed perturbations together with scRNAseq, such as Perturb-seq and CROP-seq, help uncover how cells respond to diverse stimuli or interventions (*3–5*). These approaches advance our understanding of cellular functions and regulatory networks, providing crucial insights for drug development and disease treatment (*6*). However, current experimental systems are largely optimized for cell lines, with limited applicability to primary or in vivo systems. Furthermore, combinatorial multiplexing experiments remain prohibitively expensive and technically challenging to scale (*7, 8*).

In this context, in silico approaches that predict post-perturbation cell states across diverse environments and conditions present a promising, scalable alternative and provide a path to building a virtual cell (*9, 10*). Early approaches focused on predicting cellular responses in cell types absent from experimental training data or responses to combinations of previously tested perturbations (*11–13*). However, these methods could not be generalized to unseen perturbations. Recent advances have incorporated graph-based methods and foundational models that take advantage of prior knowledge of gene expression relationships, achieving significant improvements in generalizability (*14–17*).

Among them, single-cell foundation models represent a pivotal breakthrough. By leveraging vast datasets and pre-trained transformer architectures (*18*), these models learn robust, generalizable relationships between genes and have demonstrated potential across diverse downstream tasks, including transcriptome prediction following perturbations after models are fine-tuned on specific datasets (*19*). Despite their promise, the field lacks a comprehensive framework to evaluate their capabilities systematically. Compared to tasks such as cell annotation and batch correction, perturbation prediction offers experimentally generated model input/output with well-defined objectives, making it a more robust benchmark for assessing foundation models (*15, 20, 21*). Recent studies have provided initial benchmarks comparing foundation models, other deep learning architectures, and simple baselines for perturbation effect prediction, and reported that complex models do not consistently outperform the simpler alternatives (*22–26*). While this work provides valuable insights, its evaluation was limited in dataset diversity and task coverage, leaving the question of whether such conclusions generalize across broader biological contexts.

At the same time, the Virtual Cell Challenge (*27*) has emerged as a community benchmark for perturbation prediction, attracting wide participation and fostering the development of new evaluation frameworks such as Cell-Eval (*28*). Together with the release of large-scale datasets like Tahoe-100M (*29*), these initiatives reflect a growing momentum toward standardized, transparent, and multi-metric benchmarking in this rapidly evolving field.

Motivated by these developments, we present a comprehensive benchmark that significantly expands the scope to a more diverse set of datasets, perturbation modalities, and prediction tasks. In addition to systematically evaluating model performance, we summarize common patterns across datasets and identify key biological and technical factors influencing predictive accuracy—most notably, that the magnitude of the perturbation effect strongly impacts both generalization and double perturbation prediction performance, and that in cell-type transfer settings, greater transcriptomic distance between source and target cell types is associated with reduced accuracy.

## Results

### Benchmarking models and experimental design

We comprehensively benchmarked 13 single-cell perturbation response prediction models across a broad and diverse dataset collection, alongside a suite of baselines for different tasks. Our goal was to assess how well different model architectures generalize across biological contexts, perturbation types, and prediction tasks (fig. S1). Our dataset collection spans a broad range of biological contexts, perturbation modalities, and effect sizes. It integrates large-scale public resources, recently released high-throughput perturbation screens, and in-house generated datasets. The collection covers genetic perturbations (e.g., CRISPR knockouts and activations), small-molecule treatments (e.g., kinase inhibitors, anti-aging compounds), and immune stimuli (e.g., LPS, cytokines), measured across cancer cell lines (e.g., K562, A549), pluripotent stem cells (e.g., hESCs), and primary immune populations (e.g., T cells, NK cells, monocytes), as well as non-human immune cells (e.g., mouse, rat). Of note, we incorporated datasets that include multi-cell-type profiling from the Virtual Cell Challenge, extensive drug screens across dozens of cell lines, and primary immune cell responses to cytokines. In addition, we generated two new datasets that capture the effects of drugs on both myeloid and lymphoid subsets, enabling a more systematic evaluation of cell-type transfer (Fig. 1A).

**Fig. 1.**
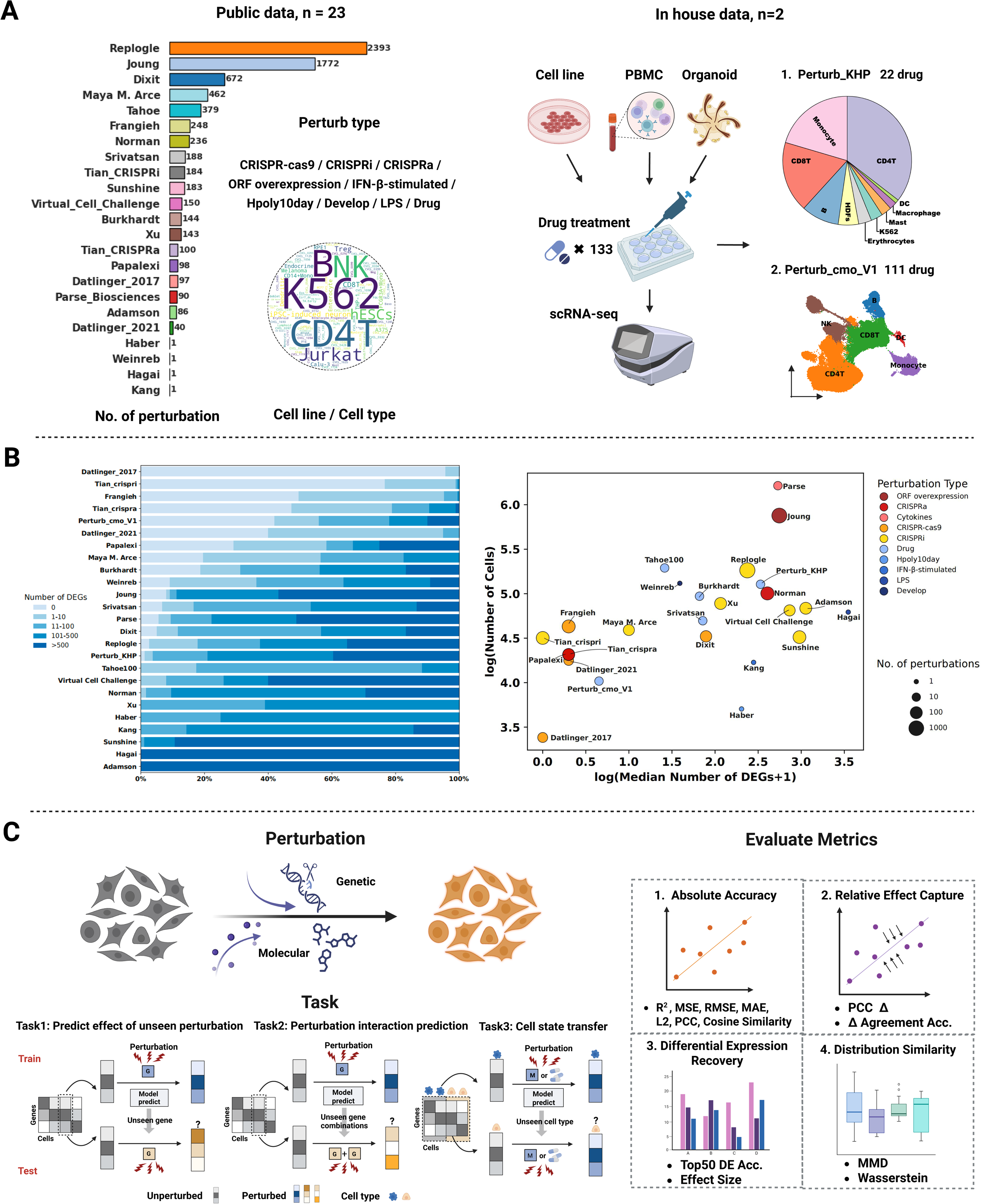
Benchmarking datasets, perturbation properties, and experimental design. **(A)** Data overview. The benchmarking collection integrates 23 public datasets and 2 in-house datasets. Public resources include large-scale CRISPR screens, drug perturbation atlases, and immune stimulation studies (numbers shown in the figure denote the total number of perturbations in the original datasets). The in-house datasets profile the effects of 133 drug treatments across lymphoid immune subsets, enabling systematic evaluation of cross–cell-type generalization. Together, these datasets cover a wide range of perturbation types (CRISPR–Cas9, CRISPRi, CRISPRa, ORF overexpression, Drug, IFN-*β*, Hpoly, LPS) across diverse cellular systems, including cancer cell lines, pluripotent stem cells, PBMCs, organoids, and non-human immune cells. **(B)** Perturbation effect size and dataset properties. Left: distribution of dataset components by percentage of differentially expressed genes (DEGs) per perturbation compared to controls. Right: scatterplot of the median number of DEGs per condition (x-axis) versus the log number of cells (y-axis), summarizing the number of DEGs across all perturbations in each dataset (including both training and testing conditions). Point size indicates the number of perturbations, and color encodes perturbation type, ranging from red (strongest transcriptional effects) to blue (weakest effects). **(C)** Benchmarking tasks and evaluation strategy. Both genetic and molecular perturbations are used in benchmarking. Models were trained on observed per-turbation–response pairs and evaluated on held-out conditions. Three tasks were benchmarked: Task 1, prediction of responses to unseen perturbations; Task 2, prediction of combinatorial perturbation interactions; and Task 3, transfer of responses across unseen cell types. Performance was assessed using multiple complementary metrics to assess absolute accuracy, relative effect capture, differential expression recovery, and distributional similarity. Created in BioRender. Tian, L. (2026) https://BioRender.com/me28fg3.

We systematically examined perturbation modalities, numbers of differentially expressed genes, E-distances, and sample sizes both across experiments and within individual experiments. This analysis revealed broad dispersion in the number of perturbations, cell numbers, and perturbation effect sizes, with substantial heterogeneity not only between studies but also among conditions within the same study. While perturbation modality may contribute to transcriptional response variation, well-designed, well-powered datasets—such as the Replogle genome-scale screen in RPE1 cells and the curated Virtual Cell Challenge—can exhibit strong signals (often > 500 DEGs for many conditions) (Fig. 1B). Beyond modality, cell-type sensitivity further modulates measurable effects, with certain immune subsets showing muted transcriptional responses to specific perturbations compared with other cell types. A full characterization of these properties and details of all datasets utilized are summarized in Table 1 and Table S1. This heterogeneity in effect size and response patterns enables a more comprehensive and discriminative assessment of model generalization across diverse biological scenarios.

**Table 1.**
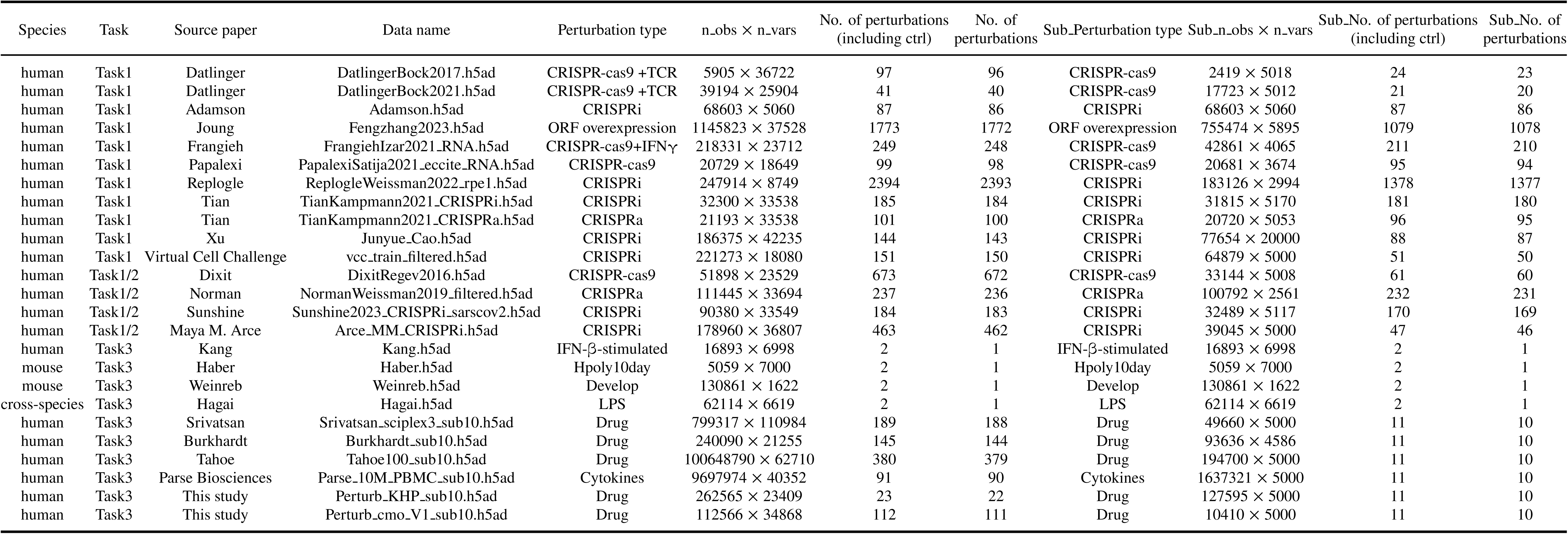
Overview of datasets used in this study.

In terms of the benchmarking object, we categorized them into three tasks focusing on their ability in different scenarios, each representing a different aspect of generalization: predicting unseen perturbations, combinatorial perturbation interactions, and responses across unseen cell types. These tasks reflect the varying complexity of perturbation prediction and the diverse experimental contexts in which these models are applied. The evaluation process is shown in Fig. 1C, where the input and output for the training and testing phases are outlined. Specifically, we focus on the following.

1. **Unseen Perturbation Prediction** (Fig. 1C, bottom left): In this task, the models were trained to predict the effects of unseen single-gene perturbations within the same cell type or cell line. Using training data that included both unperturbed and known single-gene perturbation responses, the models were tasked with inferring gene expression changes under previously unseen perturbations.
2. **Combinatorial Perturbation Prediction** (Fig. 1C, bottom middle): This task introduced additional complexity by requiring the models to predict the interaction effects of combinatorial perturbations. Based on single-gene perturbation data, models were expected to predict how new combinations of gene perturbations would influence gene expression in the same cell type or cell line.
3. **Cell state transfer** (Fig. 1C, bottom right): This task assessed the models’ ability to generalize the effect of a known perturbation across different cell types or other categories. The models were trained on unperturbed and perturbed data from various cell types to learn gene expression patterns under specific perturbations. They were then tested on previously unseen cell types, predicting their responses to the same perturbations.

To evaluate the models in the three tasks, we selected 13 representative models categorized into 4 types based on their design and model structure. (1) Foundation models (scGPT, scFoundation), pre-trained on large datasets. (2) Gene regulatory network–based models (GEARS, scELMo), which use regulatory information to predict perturbation effects. (3) Variational Autoencoder(VAE)-base model (PerturbNet, scGen, trVAE, scVIDR, scPRAM). (4) Other deep learning-based models (CPA, scPreGAN, BioLord, Squidiff). Additionally, we established a range of task-specific baselines for comparison, including simple statistical baselines (context-mean and perturbation-mean predictors) and model-based baselines (linear regression and MLP-based neural network models). Detailed baseline descriptions are provided in the Methods section (Baseline Models), and model descriptions are provided in Table S2.

Next, we implemented a standardized, multi-dimensional evaluation pipeline designed to capture complementary aspects of model performance across diverse perturbation scenarios (Fig. 1C, right). Our framework integrates four categories of metrics that together assess absolute accuracy, relative effect capture, differential expression recovery, and distributional similarity. Absolute gene expression–level agreement was quantified using error-based measures (MSE, RMSE, MAE, and L2 loss), goodness of fit (R2), and correlation metrics (Pearson’s correlation and cosine similarity). Each of these metrics was calculated on the full gene set as well as on subsets containing the top 20, 50, or 100 genes ranked by absolute score, allowing us to assess model behavior across different levels of gene prioritization. To evaluate the ability to capture relative perturbation effects, we computed expression changes (perturbed versus control) and assessed Pearson’s correlation on these delta values, together with delta direction accuracy, which measures the proportion of genes with correctly predicted up- or down-regulation. We further evaluated the recovery of differential expression by identifying DE genes in both predicted and observed profiles, then quantifying the overlap accuracy and the Spearman correlation of effect size, defined as the number of DE genes per perturbation. Finally, to assess higher-order similarity beyond mean shifts, we calculated the Wasserstein distance and maximum mean discrepancy (MMD) between predicted and observed gene expression values. Both metrics were computed per gene by comparing the distributions of predicted and observed expression values across all cells for that gene. The per-gene distances were then aggregated to yield a single scalar that summarises overall distributional fidelity for each perturbation condition. In addition to absolute metric values, we performed a ranking-based meta-analysis in which models were ranked for each perturbation and metric, and aggregated win rates were computed to capture relative performance trends across datasets and tasks.

In summary, we systematically evaluated 13 representative methods and baseline models across three core tasks, utilizing 25 datasets and 24 distinct evaluation metrics to provide a comprehensive comparison of model performance.

### Prediction of unseen single-gene perturbations

In the first task, we assessed the ability of each model to generalize to single-gene perturbations that were not observed during training, a critical measure of whether models can extrapolate to new genetic interventions (Fig. 2A). GEARS and scELMo leverage gene regulatory priors, foundation models such as scFoundation and scGPT benefit from large-scale pretraining, and BioLord and PerturbNet integrate broad perturbation–response relationships. As baselines, we used simple statistical predictors (context-mean) and model-based approaches, including linear regression and MLPs with one-hot perturbation encodings (see Methods).

**Fig. 2.**
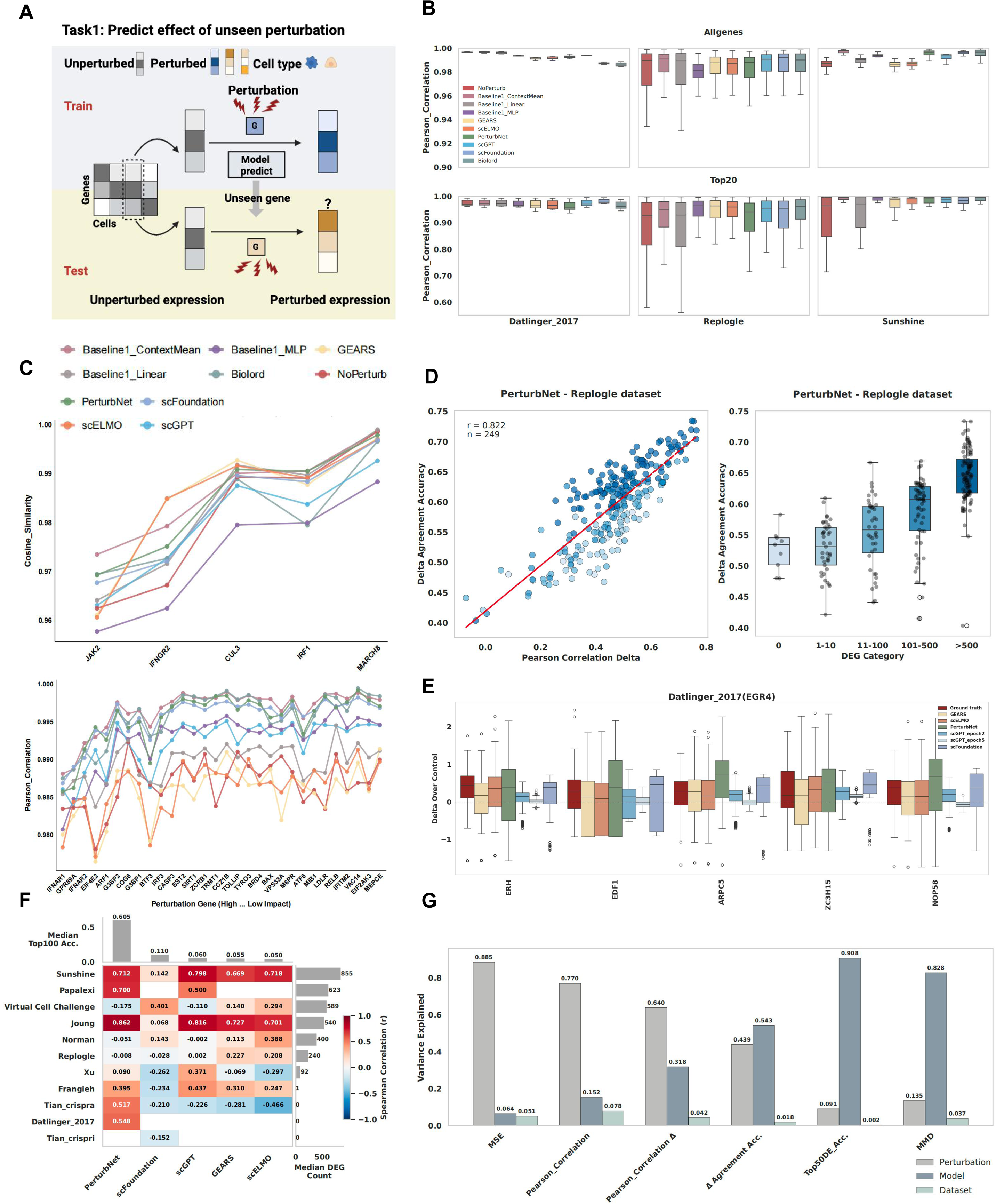
Prediction of unseen single-gene perturbations across multiple models and datasets. **(A)** Schematic overview of the single-gene perturbation prediction task. Created in BioRender. Tian, L. (2026) https://BioRender.com/ bjpu41h. **(B)** Absolute gene expression-level performance metrics. Pearson’s correlation coefficients on all genes and the top 20 genes across different datasets and models. **(C)** Cosine similarity (Papalexi dataset) and Pearson’s correlation coefficients (Sunshine dataset) on selected perturbation genes across models. The perturbation genes are ranked by impact strength. **(D)** Delta-based analysis of perturbation effect size and prediction accuracy predicted by PerturbNet model on Repologle dataset. Pearson correlation and delta direction accuracy were positively correlated (left), with larger-effect perturbations yielding higher delta direction accuracy (right). **(E)** Distribution comparison of the top 5 genes between predicted and observed expression changes on EGR4 perturbation of Datilnger 2017 dataset. **(F)** Correlation of effect sizes between true and predicted perturbations relative to control. All statistics shown are computed using the subset of Task 1 test perturbations shared by all models. Median overlap accuracy of the top 100 DEGs across datasets is shown above, and median DEG count per dataset is shown on the right. Models are ranked by overlap accuracy, datasets by median effect size. **(G)** Variance decomposition analysis showing the relative contribution of perturbation identity, model choice, and dataset identity to prediction performance across different evaluation metrics.

Using absolute gene expression–level metrics, we observed generally small performance differences between models, with Pearson’s correlation exceeding 0.95 for most datasets (Fig. 2B, fig. S2 to 7). Datasets with smaller perturbation effects tended to exhibit higher correlation-based performance and lower variance across models (e.g., Datlinger 2017 with many low-DE conditions), whereas datasets characterized by larger perturbation effects, such as Replogle and Sunshine, showed lower overall scores (Fig. 2B, fig. S3 to 7). Within datasets, effect size quantified by TRADE (transcriptome-wide impact) showed a similar trend—smaller-effect perturbations were easier to predict (Fig. 2C, fig. S8). This likely reflects the fact that small-effect perturbations are more difficult to distinguish from technical noise, whereas large-effect perturbations provide a clearer and more reproducible signal for the models to capture.

When moving to an analysis based on expression changes relative to the control, the relationship was reversed. Delta-based Pearson correlation and delta direction accuracy were strongly positively correlated, and perturbations with larger effect sizes yielded better performance (Fig. 2D, fig. S9).

Foundation models exhibited variance compression after fine-tuning, reducing the spread of predicted deltas toward a dataset-wide mean, as confirmed by higher MMD and Wasserstein distances relative to other model classes. Notably, scGPT converges to the population average and has worse distribution alignment as the training epoch increases. (Fig. 2E, fig. S10A). This “variance compression” likely arises because the distribution preservation is not in the fine-tuning objectives in the foundation model.

Evaluation of DE gene overlap accuracy and effect size ranking further supported the delta-based trends: datasets with larger perturbation effects achieved higher DE overlap accuracy and stronger concordance in effect size ranking. In this setting, PerturbNet emerged as the clear top performer, achieving a mean overlap accuracy of 0.56, whereas most other methods failed to recover DEGs reliably in several datasets (Fig. 2F, fig. S6B, C).

Variance decomposition showed that dataset and perturbation identity were primary factors for absolute and delta metrics, while DE accuracy was strongly driven by model choice, dominated by PerturbNet (Fig. 2G). To further improve interpretability, we conducted a granular variance decomposition by explicitly modeling biologically relevant attributes (e.g., KO efficiency, effect size) and technical model design factors (e.g., latent structure, loss objective). This analysis confirmed that dataset-level attributes—particularly perturbation efficiency—mainly explain variance in numerical accuracy, whereas model design choices (specifically latent space architecture) predominantly govern biological plausibility (DE accuracy) and distributional alignment (fig. S10C).

### Perturbation interaction prediction

Task 2 evaluated each model’s ability to predict gene expression responses to combinatorial perturbations. We considered three “seen levels” based on whether the individual perturbations comprising a combination had appeared in the training data: seen0 (neither perturbation seen), seen1 (one perturbation seen), and seen2 (both perturbations seen) (*14*) (Fig. 3A). Predicting seen0 combinations is most challenging, as it requires both generalization to previously unseen perturbations—analogous to Task 1—and inference of interaction effects, whereas seen2 primarily tests a model’s ability to capture the interaction effect of combinatorial perturbation. Models included are CPA, GEARS, scELMo, scFoundation, scGPT, PerturbNet, BioLord and Baseline models (see Methods).

**Fig. 3.**
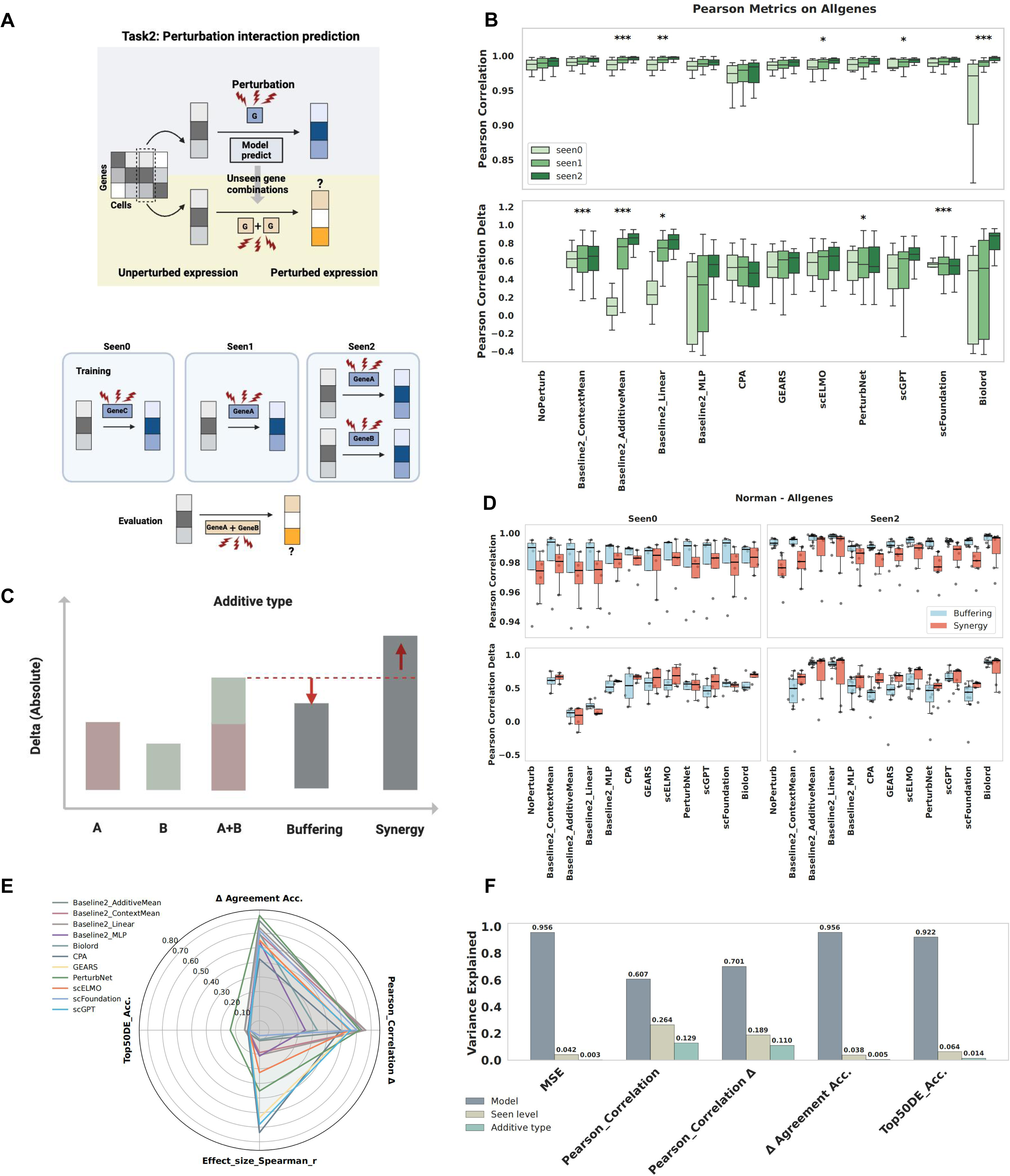
Perturbation interaction prediction performance across models and interaction types. **(A)** Schematic overview of the perturbation interaction prediction task. Bottom panel illustrates the three ”seen levels” for combinatorial perturbation evaluation: seen0 (neither individual perturbation in training data), seen1 (one perturbation in training data), and seen2 (both perturbations in training data). Created in BioRender. Tian, L. (2026) https://BioRender.com/bjpu41h. **(B)** Performance metrics for Pearson’s correlation coefficients on expressions and delta expressions across models comparing seen0, seen1, and seen2 conditions. Significance is shown. **(C)** Classification of combinatorial perturbations into buffering (combination delta smaller than additive expectation) and synergistic (combination delta larger than additive expectation) interaction types. **(D)** Comparison of Pearson correlation and Pearson correlation on deltas between buffering and synergistic combinations under seen0 and seen2 conditions across models, evaluated on all genes in the Norman dataset. **(E)** Averaged top 50 DEG overlap accuracy, effect size correlation (Spearman r), Pearson’s correlation coefficients on delta, and delta direction accuracy across all tested models for perturbation interaction prediction. **(F)** ANOVA variance decomposition analysis results showing variance contributions from model choice and interaction type factors for different performance metrics.

Across absolute gene expression–level metrics, most models showed a consistent and significant performance increase from seen0 to seen2, indicating that prior exposure to constituent perturbations in training data improves predictive accuracy. A similar trend was observed for delta Pearson correlations in some methods (Fig. 3B, fig. S11 to 16). Besides, simple baselines proved highly competitive with complex deep learning models, which is consistent with a previous study (*25*). Specifically, additive Mean and Linear models demonstrated superior performance in seen1 and seen2 scenarios, often surpassing more complex architectures in these settings where constituent perturbations were known (Fig. 3B, fig. S11 to 16).

We further classified each combination based on its deviation from the additive expectation—comparing the observed delta for the combination to the sum of deltas for its components—into buffering (combination delta smaller than additive) and synergistic (combination delta larger than additive) categories (Fig. 3C).

Across both seen0 and seen2 scenarios, we observed a distinct metric-dependent performance inversion. Buffering combinations consistently achieved higher accuracy in absolute profile correlation (PCC) compared to synergistic ones. In contrast, for Delta metrics, synergistic combinations outperformed buffering combinations. (Fig. 3D, fig. S17 to 19).

At the delta agreement accuracy and DE evaluation levels, PerturbNet again consistently outperformed all other models, replicating its dominance in Task 1, particularly in DE overlap accuracy, where it achieved substantially higher values while most other methods struggled to identify any significant DE genes (Fig. 3E, fig. S13D).

ANOVA variance decomposition revealed that model choice was the predominant determinant across nearly all metrics, overshadowing the contributions of seen levels and additive interaction types. Specifically, variance in MSE and MMD was driven primarily by model design, marking a distinct shift from data-driven limitations. Similarly, the accuracy of downstream network retrieval (Top 50 DE accuracy) was consistently governed by these design factors (Fig. 3F, fig. S13E).

### Cell state transfer

Task 3 assessed each model’s ability to generalize known perturbation responses across distinct cell types (Fig. 4A). We evaluated six methods, including scPre-GAN, scGen, trVAE, scVIDR, scPRAM, and Squidiff, using diverse datasets in which at least two cell types shared a set of perturbations. A range of baseline methods were used for comparison (see Methods).

**Fig. 4.**
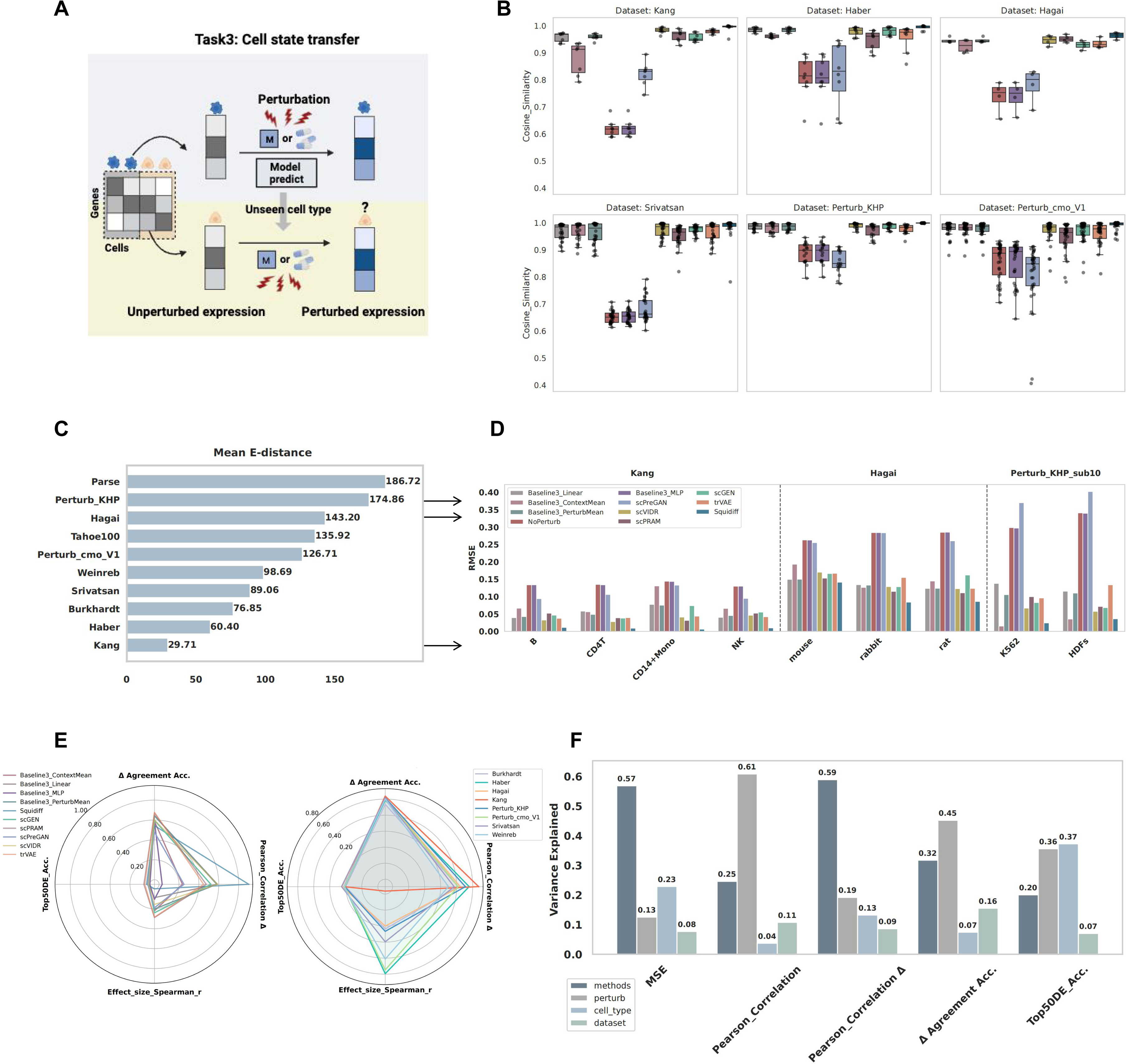
Cell state transfer performance evaluation across cell types. **(A)** Schematic overview of the cell state transfer task framework. Created in BioRender. Tian, L. (2026) https://BioRender.com/bjpu41h. **(B)** Absolute gene expression-level performance comparison across models and datasets. Cosine similarity is the evaluation metric. **(C)** The average E-distance among cell types for each dataset, ranked by values. **(D)** Model performance in predicting the responses of different cell types, evaluated using RMSE on the selected Kang, Hagai, and Perturb KHP sub10 datasets. **(E)** Averaged top 50 DEG overlap accuracy, effect size correlation (Spearman r), Pearson’s correlation coefficients on delta expressions, and delta direction accuracy across all tested models for cell state transfer prediction. **(F)** Variance partitioning analysis showing the relative contributions of model choice, perturbation identity, and cell type to overall transfer performance variability.

For absolute gene expression–level evaluation, most models (except scPre-GAN) achieved similarly high performance across datasets (Fig. 4B, fig. S20 to 24). However, notable variation was observed across datasets: in general, datasets with smaller mean E-distances (E-distance, measuring the separability of perturbed versus control distributions in high-dimensional space, reflects perturbation effect size and aligns with signal-to-noise ratios) between source and target cell types yielded better transfer performance (Fig. 4C,D, fig. S25 to 26). Smaller E-distances generally correlated with better transfer accuracy (e.g., transfer IFN-*β* stimulation in human immune subsets compared with transfer various drug effects from epithelial to K562).

But E-distance is not fully predictive. For example, results from Burkhardt showed low transfer accuracy despite a small E-distance, likely due to heterogeneous drug effects and high within-group variability, while results from Hagai, with a large domain shift and strong transfer, benefited from conserved immune activation signatures. These results indicate that Task 3 performance is shaped not only by baseline domain differences between cell types, but also by the consistency of perturbation effects across cell types (fig. S25 to 26).

Delta-level performance mirrored absolute-level trends, with trVAE leading in direction accuracy, followed by scPRAM, and Squidiff performing best in delta Pearson correlation, followed by scVIDR (Fig. 4E). At the DE level, all models performed poorly, consistent with findings from previous tasks (Tasks 1 and 2), indicating a persistent challenge in accurately predicting differentially expressed genes (Fig. 4E).

Variance decomposition quantified a functional decoupling between numerical reconstruction and biological retrieval in cross-cell-type transfer tasks. Global alignment metrics (MSE and Delta Pearson) were predominantly determined by algorithmic design factors, specifically loss formulation and latent space structure. Conversely, the accuracy of differentially expressed gene identification (Top 50 DE Accuracy) was governed by target cell identity (Fig. 4F, fig. S26B).

## Discussion

This study benchmarks single-cell perturbation response prediction across three complementary settings: unseen single perturbations, combinatorial perturbations, and cross-cell-state transfer. Across these tasks, a consistent pattern emerged. Many models achieved high accuracy when evaluated by absolute gene-expression reconstruction, yet this apparent success did not consistently translate into accurate recovery of perturbation-induced changes, differentially expressed genes, distributional similarity, or transfer across cellular contexts. Thus, the central practical message from our benchmark is that perturbation prediction performance cannot be summarized by a single global score; each metric must be interpreted in terms of the biological signal it is able to measure.

Perturbation effect size was a major determinant of this discrepancy. For weak perturbations, absolute expression metrics such as correlation or mean-squared error often yielded high scores because perturbed profiles remained close to control states and most genes were unchanged. In this regime, raw profile metrics have limited discriminative power: a model can appear accurate by reconstructing the basal expression state while failing to recover the perturbation-specific response. In contrast, delta-based metrics emphasize expression changes relative to control and become more informative when perturbation effects are large enough to rise above technical and sampling noise. DE-gene recovery imposes an even stricter requirement, because models must predict not only the correct direction of change but also sufficiently calibrated effect sizes and dispersion to support statistical detection. These observations are consistent with recent work showing that standard perturbation metrics can overestimate biological prediction accuracy and that metric design remains a critical challenge for virtual-cell benchmarking (*26, 30, 31*).

To formalize why different metrics diverge and why model rankings shift across evaluation families, we introduced a signal-decomposition framework (Fig. 5). We represent a perturbed expression profile as containing several evaluable components: the basal cellular state of the context (*B_c_*), perturbation-specific effects (*S_p_*), non-additive interaction residuals in combinatorial perturbations (*I_p_*_1,*p*2_), context-dependent modulation of perturbation responses (*C_c_*_,*p*_), and residual noise or unexplained variation (*ε*). This decomposition is intended as a conceptual evaluative scaffold, not as a mechanistic linear model of gene regulation. To quantify how much variance each component contributes to a given metric, we used linear model fitting as a statistical tool. The analysis showed that raw expression-level metrics are dominated by *B_c_*, whereas centered, delta-based, DE-based, and distributional metrics expose additional contributions from perturbation-specific, interaction-specific, and context-dependent components. This explains why model rankings can change substantially across metric families: different metrics are not redundant summaries of the same prediction problem, but probes of different biological signals.

**Fig. 5.**
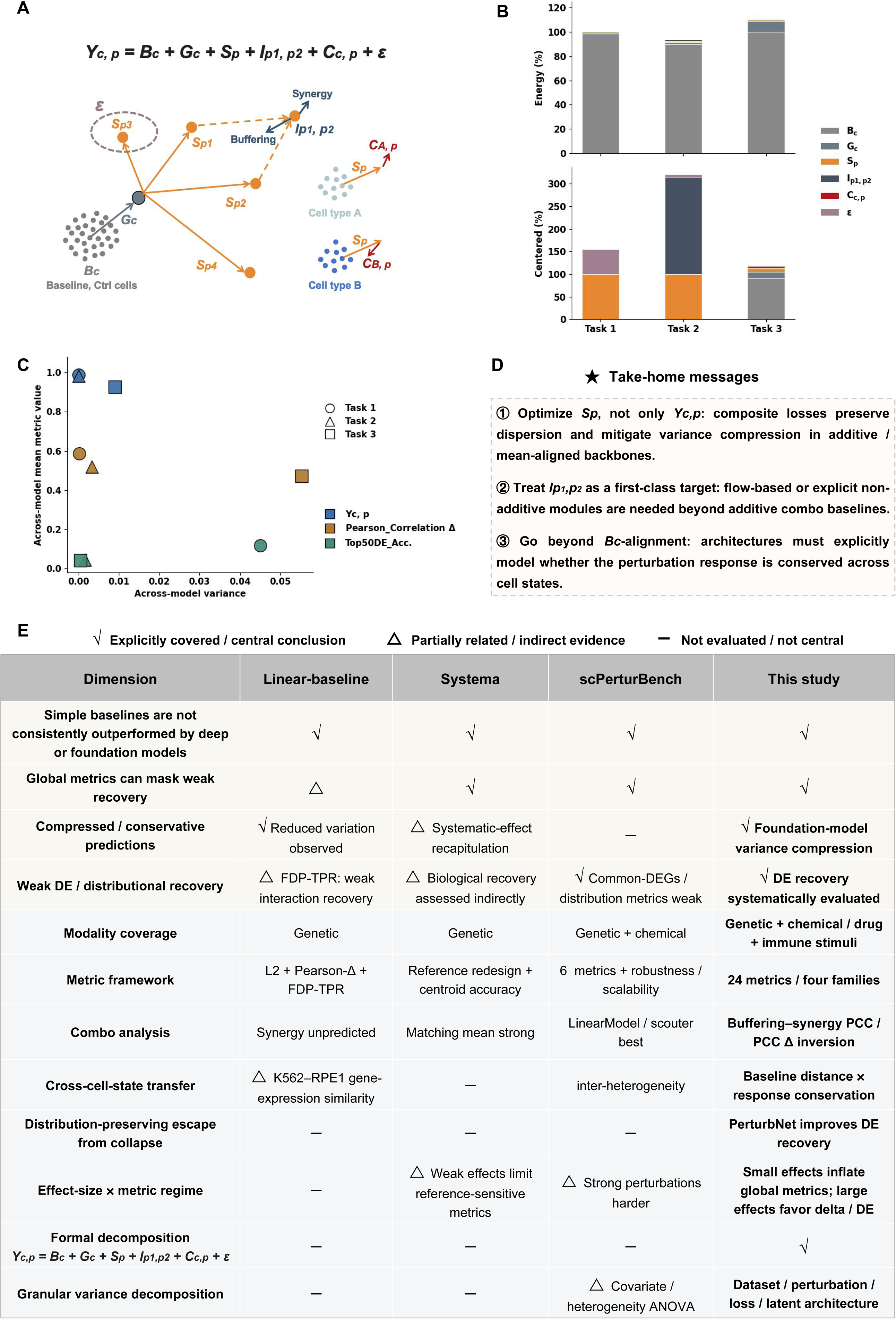
Decomposition-based interpretation of perturbation-prediction benchmarks. **(A)** Schematic of the proposed decomposition*Y_c_*_,*p*_ = *B_c_* +*G_c_* +*S_p_* +*I_p_*_1,*p*2_ + *C_c_*_,*p*_ + *ε*, separating baseline cell-state expression, systematic shifts, perturbation-specific effects, combinatorial interactions, cell-state-specific residuals of the per-

This decomposition clarifies why high reconstruction accuracy and poor DE-gene recovery can coexist. Correct DE recovery requires calibrated mean shifts and cell-to-cell variability. A model that produces overly smooth predictions may preserve global profile similarity while distorting the dispersion structure needed for reliable differential expression testing. In our benchmark, this issue was particularly evident for reconstruction-oriented AE/VAE-based models and fine-tuned foundation models, which often aligned broad expression profiles but compressed response heterogeneity. In contrast, distribution-aware generative approaches are better aligned with the single-cell nature of the task because they aim to predict a population of perturbed cells rather than only a single average profile. Consistent with this view, PerturbNet showed improved DE-gene recovery in our benchmark (*32*). However, our conclusion is not that one architecture universally solves perturbation prediction. Rather, models should be evaluated for distributional calibration, variance preservation, and DE validity in addition to mean-response accuracy.

For combinatorial perturbations, the signal-decomposition view reframes the task from predicting the full double-perturbation profile to recovering the non-additive residual beyond single-perturbation superposition. The discrepancy between buffering and synergistic combinations under different metrics should therefore be interpreted through *I_p_*_1,*p*2_ rather than through raw profile similarity alone. Buffering combinations can appear favorable under absolute profile metrics because their responses remain closer to additive or control-like expectations, whereas synergistic combinations are more visible under delta-based metrics due to larger response magnitude. Our effect-size-controlled analysis further shows that combinatorial benchmarks must separate response magnitude from interaction class before attributing performance differences to biological non-additivity. This finding extends recent observations that simple additive or linear baselines remain difficult to outperform in double-perturbation prediction (*25, 31*).

Cross-cell-state transfer revealed another distinct failure mode. Model performance was not determined by basal-state distance alone. Although large distances between control cell states can make transfer more difficult, our results indicate that transferability depends jointly on basal-state similarity and conservation of the perturbation response program across contexts. This explains why models can transfer across apparently distant contexts when the underlying response program is conserved, yet fail between closer contexts when perturbation responses are rewired. In other words, the challenge in Task 3 is not simply statistical alignment between cell types, but biological transportability of the perturbation mechanism. These task-specific diagnoses connect directly to three principles for future model development and evaluation. First, models should be trained and evaluated against perturbation-specific effects (*S_p_*) rather than raw profiles alone. Absolute expression reconstruction remains useful as a sanity check, but it should be accompanied by effect-size-stratified, centered, delta-based, DE-based, and distributional metrics. Second, combinatorial prediction should explicitly target non-additive interaction residuals (*I_p_*_1,*p*2_) beyond additive single-perturbation expectations. This will require architectures or objectives that can represent non-linear interaction structure rather than defaulting to additive superposition. Third, cross-context prediction should move beyond basal-state alignment and explicitly model whether perturbation response programs are conserved or rewired across cellular states (*C_c_*_,*p*_). To facilitate the practical application of our findings, we provide a decision framework to guide method selection based on experimental objectives and dataset characteristics (fig. S27).

Several limitations of the present benchmark also point toward concrete experimental design requirements. The datasets included in this benchmark were generated using different technologies, perturbation modalities, cell types, and experimental protocols. These differences introduce batch effects and data-quality variation that can influence both model training and evaluation. Our variance analysis indicates that intrinsic dataset properties can contribute as much to performance variation as model choice, emphasizing the need for standardized perturbation datasets with matched controls, sufficient replicates, and carefully measured perturbation efficiency. In addition, current single-cell sequencing is destructive, so we cannot observe the same cell before and after perturbation. As a result, all models infer population-level counterfactuals rather than true paired single-cell trajectories. The lack of dense temporal sampling further limits our ability to distinguish direct perturbation effects from downstream adaptation or state transitions.

Future perturbation benchmarks should therefore be designed around the unresolved signal components identified here. Multi-timepoint data would help separate immediate perturbation responses from later regulatory adaptation. Multimodal measurements, including chromatin accessibility, protein abundance, genomic variation, and molecular perturbation representations, could identify mechanisms invisible from transcriptomes alone (*33,34*). Spatial perturbation technologies may extend virtual-cell modeling from isolated cells to tissue-level response prediction (*35*). Critically, future cross-context benchmarks should deliberately sample four regimes: similar basal states with conserved responses, distant basal states with conserved responses, similar basal states with divergent responses, and distant basal states with divergent responses. Such designs would distinguish easy transfer settings from hidden context-specific failures and genuinely hard out-of-distribution regimes. These data types should not be added merely for scale, but to improve recovery of perturbation-specific effects, non-additive interactions, and context-dependent response programs. In this sense, the path toward virtual-cell models is not simply to train larger foundation models, but to build perturbation-aware systems that combine robust cellular state representations, causal perturbation data, distributional response modeling, and experimentally grounded evaluation (*36*).

In summary, our benchmark shows that current perturbation predictors often succeed on basal-state reconstruction but remain limited on the biologically decisive components of the task: perturbation-specific effects, non-additive interaction residuals, calibrated response distributions, and context-dependent response conservation. By decomposing prediction performance into these components, our study provides a framework for interpreting existing benchmarks and designing future models, metrics, and experiments for single-cell perturbation response prediction.

## Materials and Methods

### Ethics statement

Human dermal fibroblasts (HDFs) and peripheral blood mononuclear cells (PBMCs) were collected from two male donors with informed consent. The use of both HDFs and PBMCs in this study was approved by the Westlake University Ethics Committee (approval number: 20240909LXD001).

### Cell culture and drug treatment

#### Perturb KHP dataset

HDFs were cultured in DMEM medium (C11965500BT, Gibco) supplemented with 10% fetal bovine serum (FBS; C04002, VivaCell), 1% non-essential amino acids (11140-050, Gibco), 1 mM GlutaMAX (35050-061, Gibco), 55 µM 2-mercaptoethanol (21985-023, Gibco), 1 mM sodium pyruvate (11360-070, Gibco), and 1% penicillin-streptomycin (15140-122, Gibco).

PBMCs were cultured in StemPro™-34 SFM Medium (10639-011, Gibco) supplemented with StemPro™-34 Nutrient Supplement, 1 mM GlutaMAX (35050-061, Gibco), 100 ng/mL SCF, 100 ng/mL FLT-3, 20 ng/mL IL-3, and 20 ng/mL IL-6.

K562 cells, obtained from ATCC, were cultured in RPMI 1640 medium (61870-036, Gibco) with 10% FBS and 1% penicillin-streptomycin.

HDFs from two donors were seeded into 12-well plates at a density of 1×10^5^ cells per well. PBMCs from the same donors were thawed and seeded into 96-well low-attachment plates at a density of 4×10^5^ cells per well. K562 cells were seeded at a density of 1×10^5^ cells per well in either 96-well or 12-well plates. K562 cells were rested for 24 hours, while HDFs and PBMCs were rested for 48 hours, before drugs were added to each well to reach the specified final concentrations. Half of the medium was refreshed daily. A detailed list of drugs and concentrations used for each cell type is provided in the Table S3. K562 cells were collected for single-cell RNA sequencing after 3 days of treatment, while HDFs and PBMCs were collected after 4 days.

#### Perturb cmo V1 dataset

Frozen human PBMCs from three healthy donors (aged 20–25; *Stem Cell Technologies*, #70025.1) were thawed and cultured in 96-well low-attachment plates at a seeding density of 4.5 × 10^5^ cells per well. Cells from two vials originating from two donors were mixed prior to seeding. PBMCs were maintained in RPMI 1640 medium (61870-036, *Gibco*) supplemented with 10% heat-inactivated FBS (C04002, *VivaCell*) and 1% penicillin–streptomycin (15140-122, *Gibco*). Cultures were allowed to acclimate for 24 hours before drug treatment, with half of the medium replaced daily to sustain cell viability. After four days of continuous treatment, cells were harvested for downstream single-cell RNA sequencing.

A total of 111 drugs from the Drug Repurposing Compound Custom Library (L9200, *TargetMOI*) were resuspended at 10 mM in DMSO and stored at -80°C. For PBMC treatment, drugs were diluted in culture medium to a final concentration of 1 *μ*M. Control groups received DMSO at an equivalent dilution. A detailed list of drugs used provided in the Table S4.

### Sample multiplexing for single-cell RNA sequencing

Sample multiplexing was conducted using the 3’ CellPlex Kit Set A (10X Genomics, 1000261), following the manufacturer’s instructions. In brief, cells from each well were collected and labeled with unique cell multiplexing oligonucleotides (CMOs). For the Perturb KHP dataset, equal numbers of live cells from drug-treated and untreated groups were pooled into a single sample. For HDFs and PBMCs, cells from two donors treated with the same drug were mixed before incubating with the multiplexing oligos. A total of two pooled samples were prepared for K562 cells, two for HDFs, and two for PBMCs. For the Perturb cmo V1 dataset, cells were pooled at equal viability-adjusted numbers across eight treatment groups per sample, generating a total of 16 pooled samples. All samples underwent library preparation using the 10X Chromium Next GEM Single Cell 3’ Kit v3.1 (10X Genomics, 1000123).

### Datasets and preprocessing

According to the research conducted by Peidli et al. (*8*), we acquired the Datlinger (*4,37*), Dixit (*3*), Frangieh (*38*), Papalexi (*39*), Replogle (*40*), Tian (*41*), and Srivatsan (*42*) datasets from Zenodo (access link: https://zenodo.org/records/7041849). For the Datlinger and Frangieh datasets, we curated the perturbation 2 subset, excluding perturbations related to ”stimulate” and ”IFN*γ*,” as well as ”Co-culture,” while retaining only the CRISPR-Cas9 perturbations. The Sunshine (*43*) dataset was obtained from the NCBI Gene Expression Omnibus (GEO) under accession number GSE208240, where we filtered for entries classified as ”exact match,” ”no match,” or ”scrambled pair,” and excluded entries that contained non-targeting sequences in the ”guide identity.”

We further expanded our dataset collection by including the Joung and Xu datasets directly from the authors, the Virtual Cell Challenge train dataset from https://virtualcellchallenge.org/app/datasets, the Maya M. Arce dataset from GSE278572, the Tahoe dataset from https://github.com/ArcInstitute/ arc-virtual-cell-atlas/tree/main, the Parse Biosciences dataset from https://parse-wget.s3.us-west-2.amazonaws.com/10m/Parse_10M_PBMC_ cytokines.h5ad, and the Burkhardt dataset from https://www.kaggle.com/ competitions/open-problems-single-cell-perturbations/data. Due to the large size of the Tahoe dataset, which consists of 14 plates, we implemented a specialized preprocessing pipeline. For each plate, we first retained combinations of cell name and drug with at least 1,000 cells, then randomly sampled up to 100 cells per group. Subsequently, we filtered conditions to retain only those with at least 40 unique cell types within each plate, while preserving control conditions. The 14 processed plates were then merged into a single integrated dataset. For the Srivatsan, Burkhardt, Tahoe, Parse Biosciences datasets, as well as our two in-house datasets, we employed TRADE(TRADEtools v 0.99.0) to compute transcriptome-wide impact scores for each perturbation condition. Based on these scores, we selected the top 5 and bottom 5 ranked conditions for retention (detailed in Table S5). The implementation details of TRADE are provided below.

Data normalization and log transformation were performed using the scanpy(v1.10.1) toolkit. For datasets involving gene perturbations, we implemented a sample filtering process to ensure adequate coverage and minimize noise. Specifically, we applied stratified subsampling to groups with large sample sizes, limiting the maximum sample count per group to 3,000. Groups with fewer than 50 samples were excluded to reduce potential errors arising from small sample sizes. To enhance the robustness of feature selection, we calculated and retained the top 5,000 highly variable genes. Within datasets containing gene perturbations, we identified perturbed genes based on the ”condition” field and extracted their names, thereby creating a subset of genes directly influenced by the experimental conditions. Ultimately, we combined the identified perturbed genes with the highly variable genes to generate the final gene set utilized for data filtering, ensuring the preservation of the most informative genes associated with perturbations. For datasets not involving gene perturbations, we retained only the top 5,000 highly variable genes for filtering. However, for the Xu dataset, we extended the feature selection to the top 20,000 highly variable genes, as restricting to 5,000 genes resulted in insufficient perturbation conditions for analysis.

Additionally, we downloaded the Norman (*44*) and Adamson (*45*) dataset from the following links: https://dataverse.harvard.edu/api/access/datafile/6154020, https://dataverse.harvard.edu/api/access/datafile/6154417, along with the Kang (*46*), Haber (*47*), and Hagai (*48*) datasets from the scGen study (https://github.com/theislab/scgen-reproducibility/blob/ master/code/DataDownloader.py). The Weinreb (*49*) dataset was sourced from the CellOT study (*50*) (https://www.research-collection.ethz.ch/ handle/20.500.11850/609681). Given that these datasets were appropriately preprocessed in their original studies, no further data processing steps were applied. All data analysis was performed on an in-house Linux server with Ubuntu operating system (version 22.04) and the Data Science Platform of Guangzhou National Laboratory (*51*).

The DE genes are calculated using the rank genes groups function in scanpy with method=‘wilcoxon’. For each perturbation condition, cells from both the perturbation and control (‘ctrl’) groups are extracted to form a subset using normalized expression data. After obtaining the DE results, genes are sorted by the absolute values of their scores in descending order to prioritize genes with the strongest differential expression regardless of up- or down-regulation direction. The top 20, 50, and 100 DE genes based on absolute scores are then selected and used to subset the gene expression matrix for calculating the evaluation metrics.

### Models selection and setup

In this study, we selected and applied a range of advanced modeling approaches to systematically predict and compare single-cell perturbation responses. The models span diverse theoretical frameworks and algorithmic structures, ranging from baseline models to foundation models, GRN-based, VAE-based, and other deep learning-based approaches. These approaches offer strong adaptability and generalization capabilities, allowing effective application across diverse perturbation contexts through training or fine-tuning.

#### Baseline model

In single-cell perturbation prediction tasks, baseline models provide reference performance levels that facilitate the evaluation of more expressive predictive models (*28, 52–54*). Accordingly, we design task-specific baseline methods for each task, including MLP-based baselines, simple statistical predictors, and linear regression models.

### Baseline1 MLP

The baseline employs one-hot encoding to represent the cell’s gene expression under perturbation A as *p_A_*. The one-hot encoding of perturbations A is embedded into *z_A_*. The gene expression of the cell under the control (no perturbation) condition is embedded into *z*_ctrl_. The gene expression after combined perturbations can be represented as *x*^′^ = MLP(*z_A_* + *z*_ctrl_).

### Baseline1 ContextMean

This baseline predicts the average gene expression across all training perturbations for the given cell type. Let *P*_train_ denote the set of training perturbations. The prediction is defined as:

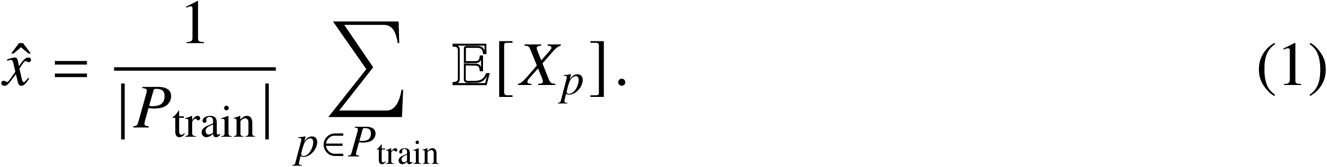

### Baseline1 Linear

This baseline uses a multi-output linear regression model that maps one-hot encoded perturbations to gene expression profiles:

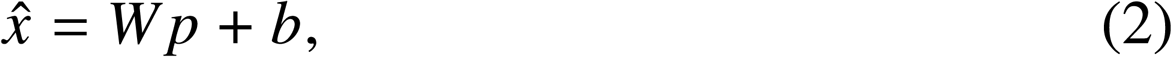

where *p* denotes the perturbation encoding, and *W* and *b* are learned parameters.

### Baseline2 MLP

We still use one-hot encoding to represent the gene expression of the cell under perturbation A as *p_A_* and perturbation B as *p_B_*. And the one-hot encoding of perturbations A and B is embedded into *z_A_* and *z_B_*, respectively. The gene expression after combined perturbations can be represented as *x*^′^ = MLP(*z_A_* + *z_B_* + *z*_ctrl_).

### Baseline2 ContextMean

This baseline is defined analogously to Task1 and predicts the average gene expression across all training perturbations for the given cell type.

### Baseline2 AdditiveMean

This perturbation-mean baseline assumes additive effects of individual perturbations. For a combination of perturbations *A* and *B*, the prediction is:

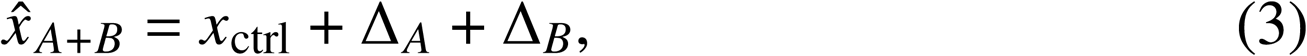

where

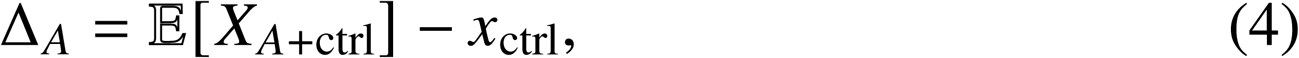

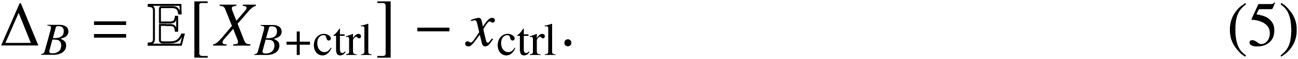

### Baseline2 Linear

This baseline uses a multi-output linear regression model with a one-hot encoding that jointly indicates both perturbed genes:

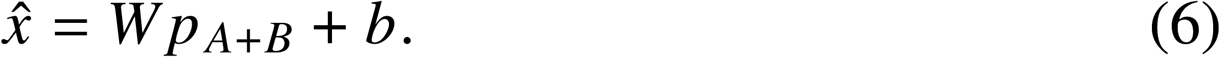

### Baseline3 MLP

We train the model for other cells under a single perturbation *A*, which is defined as *F_A_* (*Z*_other___ctrl_ + *p_A_*), where *z*_other___ctrl_ is the embedding of the gene expression of other cells under the control (no perturbation) condition, and *p_A_* is the one-hot encoding of perturbation A. For the gene expression of the target cell under perturbation A, it can be represented as *F_A_* (*z*_target___ctrl_ + *p_A_*), where *z*_target___ctrl_ is the embedding of the gene expression of the target cell under the control condition.

### Baseline3 ContextMean

This baseline predicts the average expression of the target cell type across all perturbations observed for that cell type in the training data:

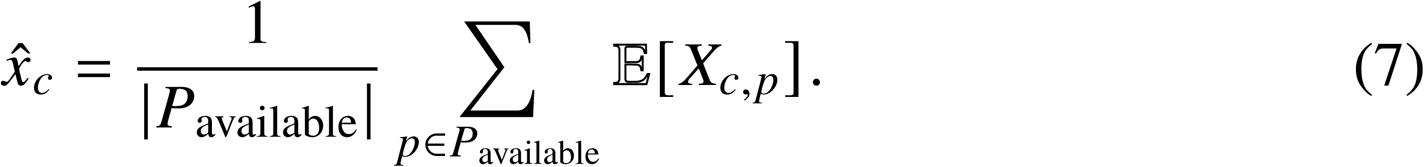

### Baseline3 PerturbMean

This baseline predicts the response of the target cell type by adding the average perturbation effect, estimated from other cell types, to the target cell type’s control expression:

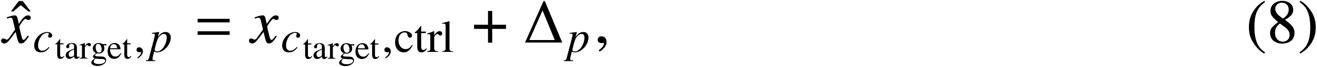

Where

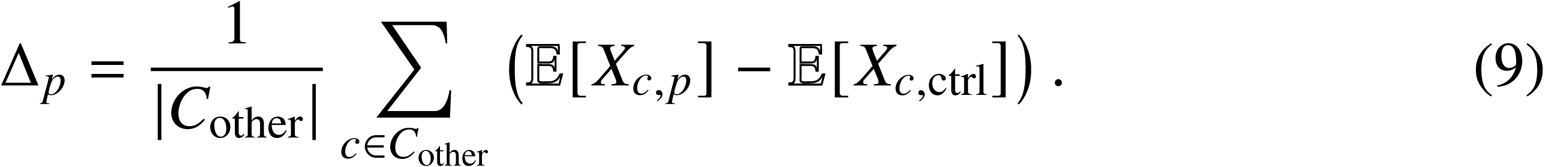

### Baseline3 Linear

This baseline uses a multi-output linear regression model that jointly encodes cell type and perturbation information via concatenated one-hot vectors.

#### Foundation model-based

The pre-trained models scGPT(v1.0) and scFoundation(v1.0) were employed for the perturbation prediction task, with fine-tuning performed based on the strategy used in the scGPT tutorial. In their approach, a Binary Condition Token was appended at each input gene position to denote the perturbation status of the gene. Besides, we implemented a strategy wherein a non-perturbed control cell was used as the input and a perturbed cell as the target. This was achieved by randomly matching each perturbed cell with a control cell, thereby forming input-target pairs. The models were trained using the default settings recommended in the tutorials, with scGPT undergoing 2 or 5 epochs of training for comparison and scFoundation trained for 2 epochs.

#### GRN-based

GEARS(v1.0) was utilized to model gene perturbations by seamlessly integrating knowledge graph information with gene embeddings. We followed the tutorial guidelines, employing the default settings as recommended.

For the scELMo(v1.0) model, we followed the tutorial’s instructions and incorporated gene embeddings derived from GPT as inputs for gene embedding in GEARS.

#### VAE-based

The VAE-based approaches, PerturbNet(v0.0.3) (*32*), scGen(v2.1.0) (*55*) trVAE(v1.0.0) (*56*), scVIDR(v1.0) (*57*) and scPRAM(0.0.2) (*58*), were utilized for single-cell RNA sequencing (scRNA-seq) perturbation prediction.

PerturbNet combines a convolutional neural network (CNN) with a variational autoencoder (VAE) to capture complex patterns in single-cell gene expression data. The model learns a latent representation of gene expression and leverages this to predict perturbation effects across different conditions. We implemented PerturbNet according to the author’s instructions, using the default architecture and hyperparameters. Note that PerturbNet requires chemical structure information (SMILES representations) as input and is therefore only applicable to drug perturbation datasets. To ensure fair comparison, main task figures present only models evaluated on shared datasets and conditions; PerturbNet’s performance on drug-specific datasets in Task 3 is presented in the Supplementary Materials.

scGen leverages a variational autoencoder (VAE) framework integrated with vector arithmetic to simulate cellular transitions between conditions, such as control and stimulation. The model encodes gene expression profiles into a latent space, reconstructing them via a decoder while learning different vectors between cellular states. This enables scGen to predict unobserved perturbation responses with high generalization. We implemented scGen following the tutorial provided by the authors, adhering to the recommended architecture and hyperparameters.

trVAE builds on the Conditional Variational Autoencoder (CVAE) framework to model perturbation effects under scenarios lacking paired control and perturbed cells. The model encodes cellular states into a shared latent space while applying condition-specific transformations via perturbation vectors. This is optimized using a masked learning objective and Maximum Mean Discrepancy (MMD) reg-ularization to align distributions across conditions. The implementation followed the author’s tutorial, with early stopping enabled and the maximum iterations set to 1000, while all other hyperparameters were kept as default.

scVIDR extends the VAE-based perturbation prediction framework by addressing cell-type-specific heterogeneity in dose-response relationships. The model employs a VAE to encode single-cell gene expression data into a latent space, then uses linear regression to predict cell-type-specific perturbation vectors ((*δ_c_*)) based on control group latent representations. For dose-response prediction, scVIDR performs log-linear interpolation on the estimated perturbation vectors to predict intermediate doses, followed by decoding back to gene expression space. The model introduces a ’pseudo-dose’ metric to characterize individual cell sensitivity to chemical perturbations by projecting cells onto the perturbation axis in latent space. We implemented scVIDR following the authors’ specifications, with default hyperparameters.

scPRAM utilizes an attention mechanism-based approach for predicting single-cell perturbation responses while accounting for cellular heterogeneity. The model consists of three main components: a VAE for encoding high-dimensional gene expression data into latent space, optimal transport based on the Sinkhorn algorithm to match unpaired cells before and after perturbation, and an attention mechanism to compute perturbation vectors for test cells. The attention mechanism treats the test cell’s latent representation as a query, using cosine similarity to identify the most relevant training cells, then computes weighted perturbation vectors through a softmax-normalized attention score. This approach enables scPRAM to capture individual cell-specific responses rather than relying on population averages. We implemented scPRAM with default parameters.

#### Other deep learning-based

The Compositional Perturbation Autoencoder (CPA,v0.8.7) (*59*) combines deep learning flexibility with linear model interpretability to predict cellular responses to perturbations. The model uses an encoder to map gene expression data into a disentangled latent space, removing perturbation-specific information via adversarial training. The latent embedding is then combined with perturbation and covariate embeddings, scaled by a learned dose-response function for cell-specific predictions. The decoder reconstructs gene expression profiles, enabling predictions for unseen perturbations. We followed the authors’ tutorial and used the default hyperparameters for implementation.

scPreGAN(v1.0) (*60*) integrates autoencoders with generative adversarial networks (GANs) to predict cellular responses to perturbations. The architecture includes an encoder, two generator networks, and two discriminators. The encoder extracts features, while the generators predict perturbed data from latent vectors. Pre-training combines reconstruction and adversarial losses, optimizing data fidelity and distribution realism. We implemented scPreGAN based on the authors’ GitHub tutorial, adhering to default settings and niter set to 1000.

Biolord(v0.03) (*61*) utilizes a deep generative framework to disentangle single-cell data into known and unknown attributes through a decomposed latent space. The model separates each known attribute (cell type, time, perturbation) into dedicated subnetworks while capturing unknown attributes in regularized embeddings. A generator module reconstructs cellular measurements from these disentangled representations. The framework is optimized using a composite loss function that balances reconstruction accuracy with information minimality constraints, enabling counterfactual predictions by modifying specific attributes while preserving cell-specific signatures. We implemented Biolord following the official tutorial with default parameters and max epochs set to 1000.

Squidiff(v1.0.8) (*62*) is a conditional diffusion-based generative framework designed to model single-cell transcriptomic dynamics under diverse biological conditions. The model integrates a semantic encoder to map gene expression profiles into a biologically meaningful latent space and a denoising diffusion implicit model (DDIM) to generate transcriptomes. By manipulating semantic latent variables via interpolation and vector arithmetic, Squidiff enables the prediction of cellular responses to perturbations across cell types. The model is trained following the official implementation with default training settings, using a diffusion process with 1000 timesteps, with input gene expression profiles restricted to the top 200 highly variable genes (HVGs)..

### Evaluation metrics

In evaluating model performance, selecting the appropriate assessment metrics is essential. To provide a comprehensive evaluation, this study employs a standardized, multi-dimensional evaluation framework that integrates four complementary categories of metrics: absolute accuracy, relative effect capture, differential expression recovery, and distributional similarity. These metrics offer a holistic view of the model’s predictive accuracy, biological interpretability, and distributional consistency across various perturbation scenarios.

Specifically, absolute accuracy is assessed through error-based measures (such as MSE, RMSE, MAE, and L2 loss), which quantify the deviations between predicted and actual gene expression values; goodness-of-fit metrics (such as R²), which evaluate the model’s ability to account for overall data variability; and correlation analysis (including Pearson correlation and cosine similarity), which measures the linear relationship and directional alignment between predicted and actual values. Relative effect capture is evaluated by computing Pearson correlation on delta values (expression changes between perturbed and control conditions) to assess the model’s ability to predict perturbation-induced changes, along with delta direction accuracy, which quantifies the proportion of genes with correctly predicted up- or down-regulation patterns. Differential expression recovery is measured through DE overlap accuracy, which assesses the agreement between predicted and observed differentially expressed genes, as well as Spearman correlation of effect sizes, which evaluates the consistency of perturbation magnitudes across different conditions. Finally, distributional similarity metrics (such as MMD and Wasserstein distance) assess the alignment between predicted and actual gene expression distributions, capturing the nuanced distributional characteristics in high-dimensional data.

#### Absolute Accuracy Metrics

**Mean Squared Error(MSE)** is a commonly used metric for evaluating model error, quantifying the mean squared difference between predicted and actual values. MSE is particularly suited to assessing prediction bias and overall accuracy, given its sensitivity to larger errors. In calculation, MSE squares the prediction error for each sample and then averages these values. The formula is as follows:

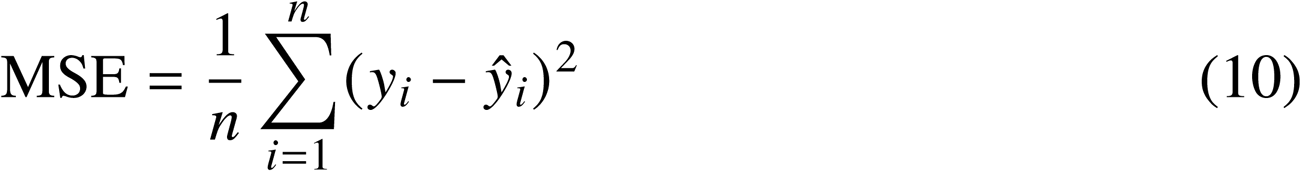

Where *y_i_* is the actual value, *ŷ_i_* is the predicted value, and *n* is the number of samples.

**Root Mean Squared Error(RMSE)** is similar to MSE, being the square root of MSE, but its units are consistent with the original data, providing a more intuitive reflection of the magnitude of prediction errors. It is obtained by taking the square root of MSE, defined as follows:

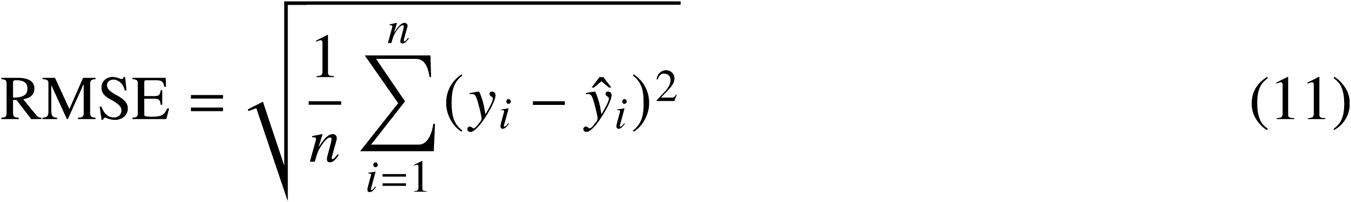

**Mean Absolute Error(MAE)** measures the absolute error between predicted and actual values, is less sensitive to outliers than MSE, and is suitable for scenarios where one does not wish to penalize large errors. MAE was calculated by taking the average of the absolute difference between the predicted and actual values, defined as follows:

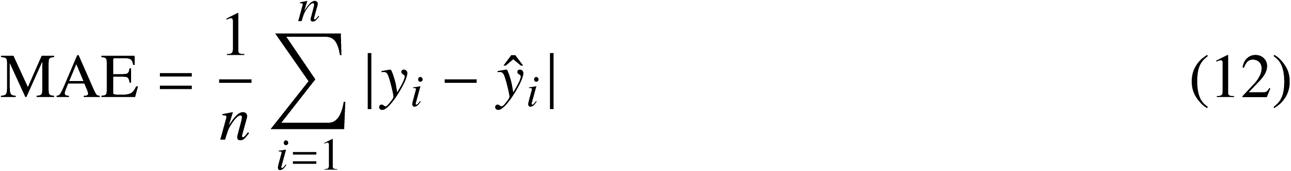

**L2 Norm(L2)** is used to measure the Euclidean distance between predicted and actual values, serving as an effective tool for describing the error space. It calculates the L2 norm of the difference vector between predicted and actual values, defined as follows:

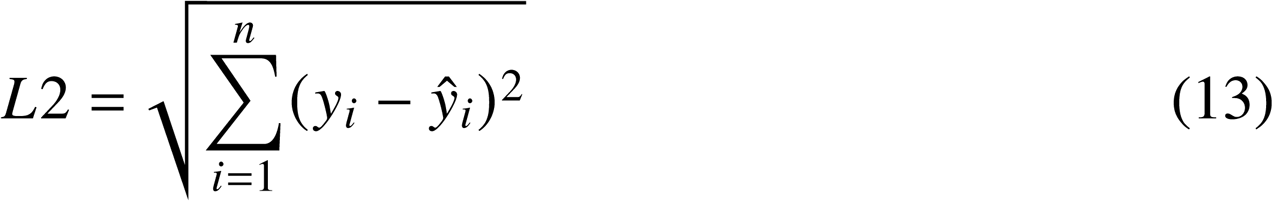

**R-squared(***R*^2^) is used to measure a model’s ability to explain the total variance, serving as an indicator of model fit. It represents the degree of correlation between predicted and actual values and is defined as:

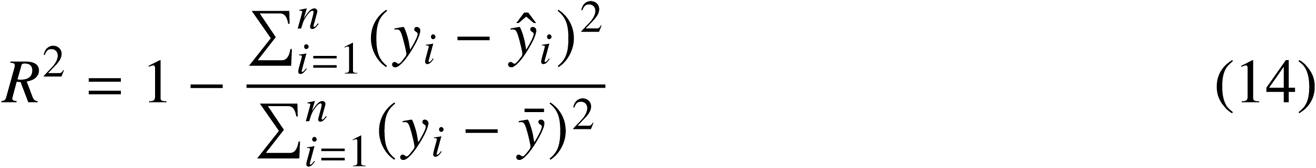

In this formula, *ȳ* is the mean of the actual values. The numerator, 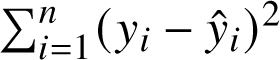, represents the residual sum of squares (RSS), which is the total squared error. The denominator, 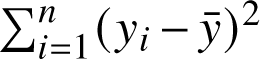, represents the total sum of squares (TSS) of the actual values.

**Pearson’s Correlation** measures the linear relationship between predicted and actual values, making it suitable for analyzing the strength of association between datasets. It is calculated by dividing the covariance between predicted and actual values by the product of their standard deviations, defined as:

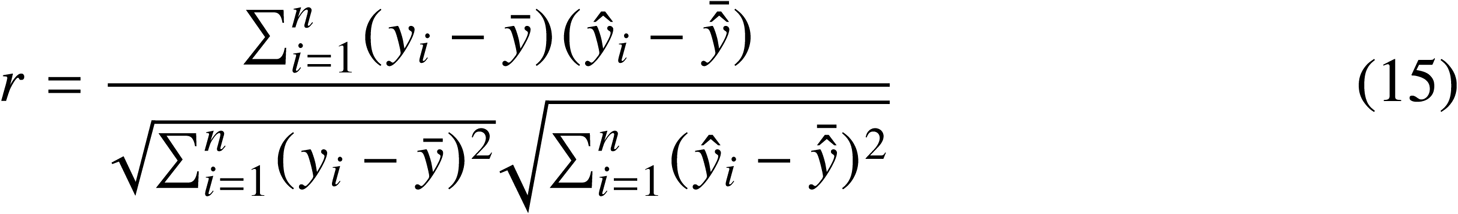

**Cosine similarity** measures the directional similarity between two vectors, making it suitable for evaluating the similarity between model predictions and actual results. The cosine similarity between the predicted and actual value vectors is defined as:

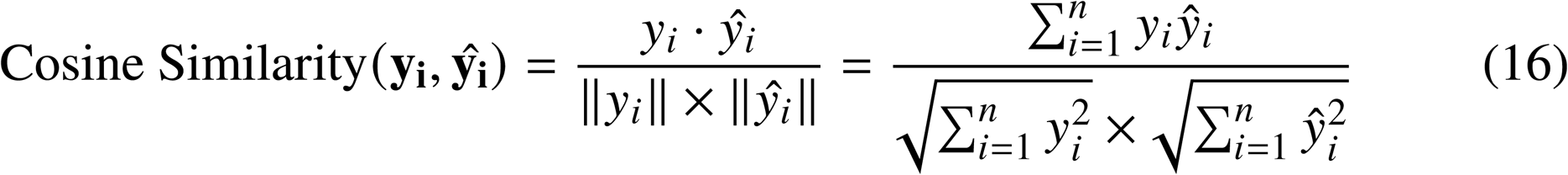

#### Relative Effect Capture Metrics

**Pearson’s Correlation** Δ measures the change in the linear relationship between the predicted and actual values when adjusted for control conditions. This metric is designed to assess how the perturbation affects the correlation between predicted and actual values relative to the control condition. Specifically, Pearson’s Correlation Δ is calculated by first subtracting the control values from both the predicted and actual data, and then computing the Pearson correlation coefficient on the adjusted data.

**Delta Direction Accuracy** evaluates a model’s ability to correctly predict the directionality of gene expression changes under perturbation conditions. This metric assesses whether genes are predicted to be upregulated or downregulated in the same direction as observed experimentally. For each perturbation condition, we first compute the expression changes relative to the control by calculating delta values for both predicted and true expression profiles. The delta direction accuracy is then defined as the proportion of genes where the predicted and observed expression changes have the same directional sign, calculated as:

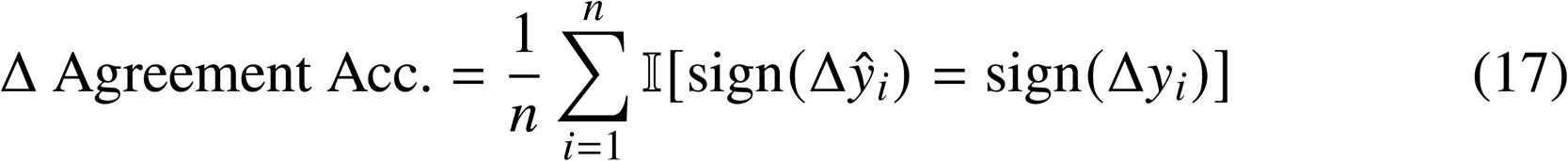

where

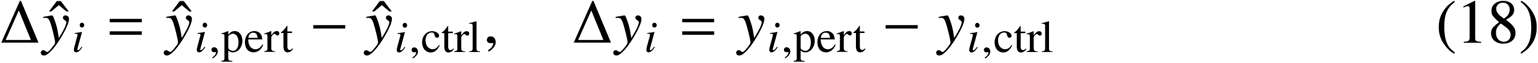

represent the predicted and true expression changes for gene *i*, respectively. Here, *ŷ*_*i*,pert_ and *y_i_*_,pert_ denote the mean predicted and true expression values under perturbation, while *y_i_*_,ctrl_ represents the mean control expression. The indicator function *I* [·] equals 1 when the condition is true and 0 otherwise, and sign(·) denotes the sign function. This metric takes values of 0 or 1, where 1 indicates perfect directional agreement and 0 indicates directional disagreement.

#### Differential Expression Recovery Metrics

**DE Overlap Accuracy** evaluates how well perturbation prediction models recover the correct differentially expressed genes compared to ground truth experimental data. This metric assesses the biological relevance of predictions by measuring the concordance between predicted and observed transcriptional responses.

For each perturbation condition, we first identify differentially expressed genes (DEGs) by comparing both predicted and true perturbed expression profiles against their respective control conditions using the Wilcoxon rank-sum test. DEGs are filtered based on adjusted p-values (Benjamini-Hochberg correction) with a significance threshold of *p_adj_* < 0.05 and ranked by absolute log fold change. The DE overlap accuracy is computed at multiple granularity levels by comparing the top-ranked DEGs between predicted and true conditions. For each top-*N* threshold (*N* ∈ 10, 20, 50, 100, 200, 500), the metric is defined as:

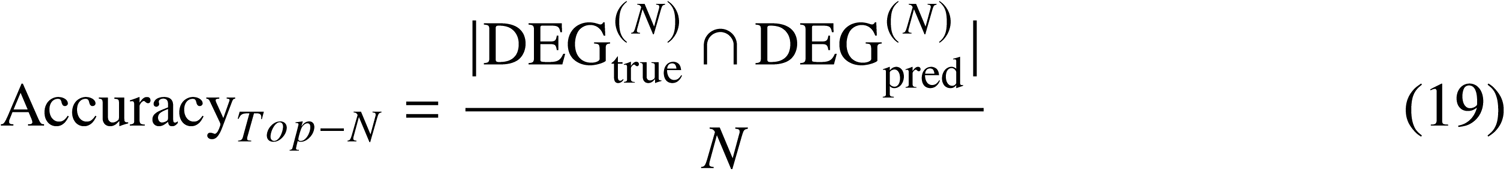

where DEGtrue^(^*^N^*^)^ and DEGpred^(^*^N^*^)^ represent the sets of top-*N* DEGs from true and predicted data, respectively, and | · | denotes set cardinality. If the number of significant DEGs in either the predicted or true condition is less than *N*, the accuracy for that *N* is not calculated. A maximum of 500 cells per condition is used, with random sampling applied when necessary. The metric ranges from 0 to 1, with 1 indicating complete overlap between predicted and true DEGs.

**Effect Size** is defined as the number of significantly differentially expressed genes (DEGs) per condition (*N*_sig_, *p_adj_* < 0.05). The distribution of effect sizes in the true datasets is examined to characterize the transcriptional response across conditions. The relationship between predicted and true effect sizes is quantified using the Spearman rank correlation:

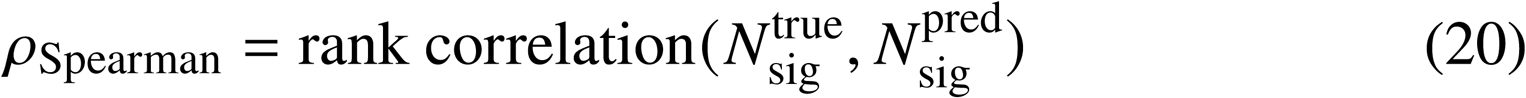

where each vector contains paired effect sizes for all conditions in a dataset. Conditions are excluded from the calculation if there are fewer than 5 valid paired observations, or if either the predicted or true values have zero variance.

#### Distributional Similarity Metrics

**Maximum Mean Discrepancy(MMD)** is used to measure the distance between two distributions, making it suitable for evaluating the difference between predicted and actual distributions. Specifically, we computed per gene (i.e., it quantifies the distributional distance between the predicted and observed expression values across all cells for that gene). After obtaining one MMD value for every gene, we then aggregate these per-gene distances to yield a single scalar that summarises overall distributional fidelity for the perturbation condition. It calculates the mean discrepancy between the predicted and actual data using a kernel function, defined as:

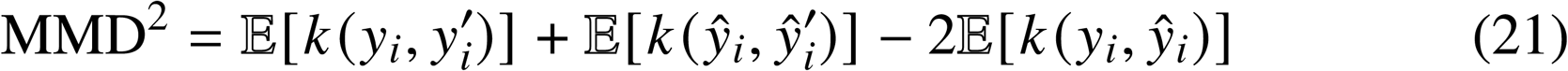

Here, *y_i_* and *ŷ_i_* represent samples from two different distributions, while *y′_i_* and *ŷ′_i_* are samples within the same distribution. *k*(*y_i_*, *ŷ_i_*) is the kernel function, and we use the RBF kernel in this study.

**Wasserstein Distance** is used to measure the geographical distance between two probability distributions and is commonly used to assess the similarity between distributions. By solving the optimal transport problem, it calculates the cumulative discrepancy between the two distributions, defined as:

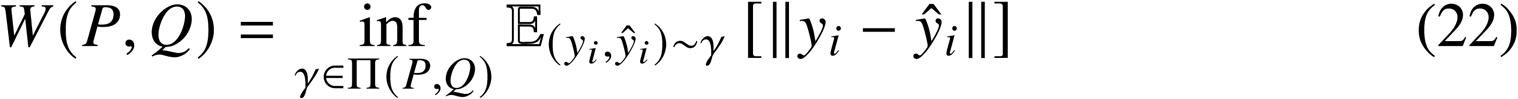

In the formula, *P* and *Q* are two different probability distributions, and *γ* ∈ Π(*P*, *Q*): represent the set of all joint distributions from *P* to *Q*.

#### Ranking-based meta-analysis

We conducted a comprehensive ranking-based meta-analysis to evaluate model performance across four standardized dimensions. The evaluation framework comprised four key dimensions, each weighted equally at 25%: (1) Absolute Accuracy, measuring prediction accuracy through error metrics (MSE, RMSE, L2, MAE) and correlation metrics (Pearson correlation, cosine similarity, R²); (2) Relative Effect Capture, assessing perturbation effect detection using Pearson correlation delta and delta agreement accuracy; (3) Differential Expression Recovery, evaluating DE gene identification capability through Top50 accuracy and Spearman correlation; and (4) Distribution Similarity, measuring distributional fidelity via Maximum Mean Discrepancy (MMD) and Wasserstein distance. For the first, second, and fourth dimensions, metrics were calculated for both all genes and the top 20 differentially expressed genes, while the third dimension used all genes only.

For score calculation, models were ranked within each metric for each task separately. Error-based metrics (MSE, RMSE, L2, MAE, MMD, Wasserstein, and execution time) were reverse-ranked, where lower values received higher normalized scores. Each dimension score was computed as the average of normalized rank scores for all constituent metrics within that dimension. The Overall Score was calculated as the unweighted average of the four dimension scores, excluding time and user-friendliness metrics, which were displayed separately for reference. Missing values were imputed using the mean value of the respective metric across all models within each task to ensure complete comparisons.

The visualization employed a multi-element approach where point size represents normalized metric values within each column (larger points indicating higher values), point color indicates rank within each column using the RdGy colormap (darker colors representing better ranks), horizontal bar charts display Overall Scores with numerical labels, and star symbols represent user-friendliness levels (1 star = good, 2 stars = better). This ranking-based meta-analysis provides task-specific model comparisons while maintaining consistency across evaluation criteria and effectively handling the heterogeneity of available metrics across different models and datasets.

### TRADE

To assess the transcriptome-wide impact of perturbations, we employed the TRADE (Transcriptome-wide Analysis of Differential Expression, TRADEtools v 0.99.0) (*63*) method, which quantifies the total effect of a perturbation across the entire transcriptome. TRADE addresses the challenges of noise and undetected effects that are common in single-cell CRISPR screens, such as Perturb-seq, by providing a robust estimation of the true differential expression effects. The analysis was performed following the tutorial provided by the authors. Differential expression analysis was carried out using DESeq2(v 1.42.1), incorporating both condition and batch information to account for potential variations between replicates. The differential expression results were then integrated into the TRADE framework, which calculates a transcriptome-wide impact score. This score reflects the global shift in gene expression distributions caused by the perturbation relative to the control condition. TRADE’s capability to summarize these effects enables the systematic ranking and comparison of perturbations across multiple datasets.

### E-distance

E-distance is a metric used to measure the difference between two distributions in high-dimensional space, defined as the mean squared Euclidean distance between samples. In this study, we employ E-distance to quantify the high-dimensional distribution differences across different datasets, utilizing scPerturb (v0.1.0) for the calculations. For samples originating from two distributions, we first compute the Euclidean distance between each pair of samples and then calculate the average distance within each distribution as well as the average distance between the two distributions. Prior to these calculations, we standardize the single-cell data, filter out low-expression cells (min counts=100) and low-expression genes (min cells=50), and select the top 2000 highly variable genes as features for analysis. Detailed methodologies can be referred to in the scPerturb (*8*) publication.

### Statistical Analysis

All statistical analyses were conducted using Python (v3.12.3) with the pandas (v1.5.3), statsmodels (v0.14.2), and NumPy (1.26.4) libraries. For each task, we performed ANOVA tests to assess the significance of performance differences across models and various experimental factors specific to each task design. Task-specific ANOVA models were implemented using ordinary least squares (OLS) regression, where all variables were treated as categorical factors using the C() notation in statsmodels.

For Task 1, we used the model:

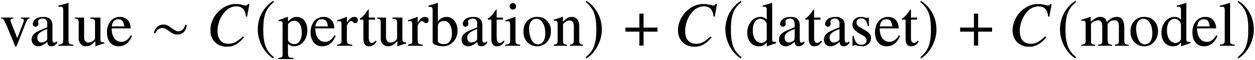

where perturbation, dataset and model served as categorical factors.

For Task 2, we used:

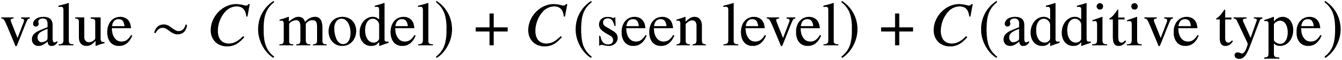

For Task 3, we used:

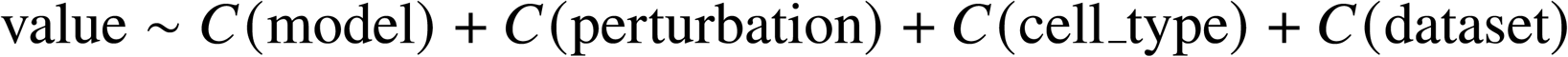

Post-hoc analysis with Tukey’s HSD test was conducted to perform pairwise comparisons between models when a significant effect (*p* < 0.05) was detected. All analyses were performed using Type II Sum of Squares for ANOVA, and results were considered statistically significant at *p* < 0.05.

In addition to the task-specific ANOVA models described above, which focus on experimental design factors, we further conducted a more granular variance decomposition analysis to better disentangle the contributions of dataset characteristics and model design choices to performance variability. For dataset-level factors, we extracted condition-specific features including: (1) sequencing depth (mean counts per cell), (2) perturbation effect size (mean absolute log2 fold change between perturbed and control cells), (3) KO/KD efficiency for genetic perturbations (Tasks 1-2), and (4) cell type heterogeneity for drug perturbations (Task 3, quantified by Shannon diversity index). For model-level factors, we characterized: (1) preprocessing strategies (normalization method, feature selection, batch correction), (2) loss function design (reconstruction, distributional, adversarial, and auxiliary objectives), and (3) latent space architecture (type, dimensionality, and structure). All continuous features were categorized into three levels (Low, Medium, High) using quantile-based binning. The granular ANOVA models included the following categorical factors. For Tasks 1 and 2, the model incorporated sequencing depth, perturbation effect size, KO/KD efficiency, preprocessing strategy, loss function design, and latent space architecture as main effects. For Task 3, the model incorporated sequencing depth, perturbation effect size, cell type heterogeneity, preprocessing strategy, loss function design, and latent space architecture as main effects. Variance proportions were calculated by dividing each factor’s sum of squares by the total sum of squares across all factors (excluding residuals).

### Response-component decomposition analysis

To interpret how different evaluation readouts reflect different sources of variation in post-perturbation expression, we performed a response-component decomposition on the observed profiles from the common model-evaluation subset of each task. For each task, the common subset was defined by retaining only the evaluated entries shared across the corresponding prediction models. Each entry was indexed by dataset and perturbation condition, with cell type additionally included when cell-state transfer was evaluated. For each entry, we computed a pseudobulk expression profile *Y_c_*_,*p*_ by averaging normalized log-transformed expression across cells assigned to cell state or cell type *c* and perturbation condition *p*.

We represented each observed post-perturbation profile using the diagnostic decomposition

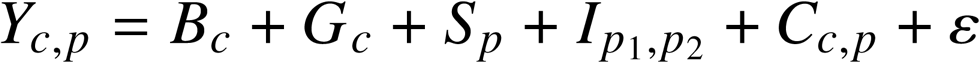

Here, *B_c_* denotes the baseline control expression of cell state *c*; *G_c_* denotes the systematic shift shared across perturbations within the same cell state; *S_p_* denotes the perturbation-specific response; *I_p_*_1,*p*2_ denotes the non-additive interaction term for combinatorial perturbations; *C_c_*_,*p*_ denotes the cell-state-specific residual of the perturbation response, corresponding to the cell-state-by-perturbation interaction term; and *ε* denotes the empirical noise floor.

For each dataset and task, *B_c_* was estimated from control cells. Perturbation-induced changes were calculated relative to *B_c_*, and the average change within each cell state was used to estimate *G_c_*. After removing *B_c_* and *G_c_*, the remaining response was treated as the perturbation-associated residual. For tasks without cell-state transfer, this residual was used as *S_p_*. For combinatorial perturbations, *I_p_*_1,*p*2_ was quantified as a separate non-additive diagnostic term, calculated as the deviation of the observed combination response from the additive expectation of the two corresponding single-perturbation responses. For cell-state transfer, the residual response was further separated into a perturbation-average component shared across cell states and a cell-state-specific residual *C_c_*_,*p*_. Components that were not applicable to a given task were set to zero. The empirical noise floor *ε* was estimated from control cells using a split-half procedure.

Component contributions were quantified as sum-of-squares energies across evaluated entries and genes. We generated two normalized summaries. In the energy view, each component was normalized by the total post-perturbation expression energy. In the centered view, expression was first centered by subtracting the gene-wise grand mean across evaluated entries, and component energies were then normalized by the centered total energy. Dataset-level fractions were averaged within each task. Because this decomposition was used as a component-wise diagnostic rather than a strictly orthogonal ANOVA partition, normalized fractions were not constrained to sum to 100%.

To relate the decomposition to model behavior, we summarized three representative evaluation readouts from the existing benchmark outputs: full-profile *Y_c_*_,*p*_, Pearson correlation Δ, and top 50 DEG overlap accuracy. For each task–readout pair, we calculated the across-model mean and sample variance across retained prediction models. NoPerturb and Baseline-related models were excluded from this analysis.

### Perturbation prediction on unseen genes

To assess the model’s response to perturbations in single genes that it has not seen, we employ a masking strategy during model training to simulate the absence of specific genes in the training dataset. Specifically, there are multiple choices for perturbed genes in each dataset, and following the same protocol as GEARS, we randomly split perturbed genes at the gene level into 75% for training and 25% for testing. During the training process, the perturbation response data for these genes are completely excluded from the training set, forcing the model to learn based solely on the perturbation features of other genes. Subsequently, the model predicts the perturbation responses for these unseen genes in the test set to evaluate its performance on unknown single-gene perturbations.

### Perturbation prediction on combinatorial perturbation

For predicting combinatorial gene perturbations, we followed the same protocol as GEARS and randomly selected multiple gene pairs from each dataset. We designed the following three training configurations to evaluate the models’ generalization capabilities under varying levels of prior knowledge: Seen0: In this configuration, neither the gene pair nor their individual single-gene perturbations appear in the training data. This setup assesses the model’s ability to predict responses to completely novel gene combinations. Seen1: The training set includes single-gene perturbation data for one gene of the pair, while the other gene remains unseen. This configuration tests whether the model can leverage partial prior information to predict the effects of combinatorial perturbations. Seen2: The training data contains single-gene perturbation data for both genes in the pair, but their combined perturbation is not included in the training stage. This setup evaluates the model’s capacity to infer combinatorial perturbation responses based on individual gene perturbation information. By randomly selecting gene pairs, we ensured that the perturbation response of each gene combination in the test set was entirely novel to the model, thereby providing a comprehensive evaluation of its generalization performance in predicting combinatorial gene perturbations.

### Perturbation prediction on unseen cell types

In the cell state transfer prediction task, we explored the models’ generalization capabilities across different cell types. We trained the models using perturbation data from multiple cell types or cell lines, to predict the perturbation responses of a target cell type that was masked during training. Specifically, the training set included perturbation response data from only a subset of cell types, while data from the target cell type were completely excluded. The models were then tasked with predicting the perturbation responses for the unseen target cell type based on the information learned from the training set. This approach allows us to evaluate the models’ applicability and robustness across different biological contexts.

## Acknowledgments

We acknowledge the support of the Data Science Platform of Guangzhou National Laboratory and the Bio-medical Big Data Operating System (Bio-OS). We thank Lead Healthcare.AI for assistance with experiments.

## Author contributions

Conceptualization: Y.Y. and L.T. Methodology: L.L., Y.Y., X.L. and L.T. Data curation: L.L., Y.Y., Y.F., Y.C. and L.T. Formal analysis: L.L. and Y.Y. Investigation: L.L., Y.Y., X.F., S.L., B.L. and W.R. Validation: L.L., Y.Y., Y.F. and L.T. Software: L.L., Y.Y., Y.F., W.L. and L.T. Resources: J.C., X.L. and L.T. Visualization: L.L., Y.Y. and Y.F. Writing—original draft: L.L., Y.Y. Y.F., W.L., X.F. and L.T. Writing—review & editing: L.L., Y.Y., Y.F., W.L, X.F., S.L., Y.C., B.L., W.R., J.K., S.Z., J.C., X.L. and L.T. Supervision: Y.Y., X.L. and L.T. Project administration: L.L., Y.Y. and L.T. Funding acquisition: L.T.

## Data, code, and materials availability

All data and code needed to evaluate and reproduce the conclusions in the paper are present in the paper and/or the Supplementary Materials. The in-house datasets and the analysis scripts generated in this study are permanently archived in Zenodo at https://doi.org/10.5281/zenodo.20823660. Except for the data unique to this study, all other datasets used are from public resources. For specific details on their usage, please refer to the ”Materials and Methods” section of the manuscript. A comprehensive list of datasets and their download sources can be found in Table 1 and Table S1. As an additional resource, the scripts are also available via GitHub at: https://github.com/TianGzlab/scPerturbBench. A website hosting the data and the visualization can be viewed at: luyitian. github.io/PerturbArena. The website is continually updated and may contain new models and show different results compared to the manuscript. This study did not generate new materials.

