## Supplementary material for "A Systematic Comparison of Single-Cell Perturbation Response Prediction Models": fig. S1

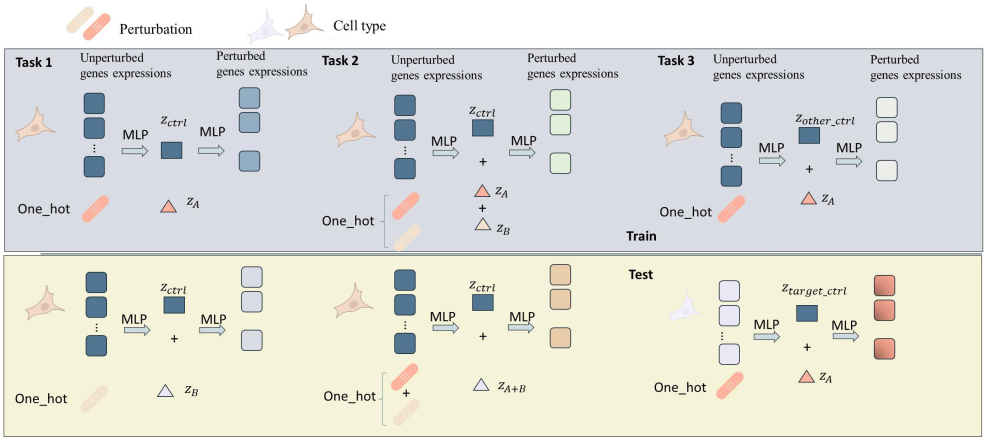


**Fig. S1. Summary of Baseline_MLP model used in this study. Created in BioRender. Tian, L. (2026) `https://BioRender.com/wwdkl0n`.**
