## Supplementary figures and images for "A Systematic Comparison of Single-Cell Perturbation Response Prediction Models"

### fig. S2

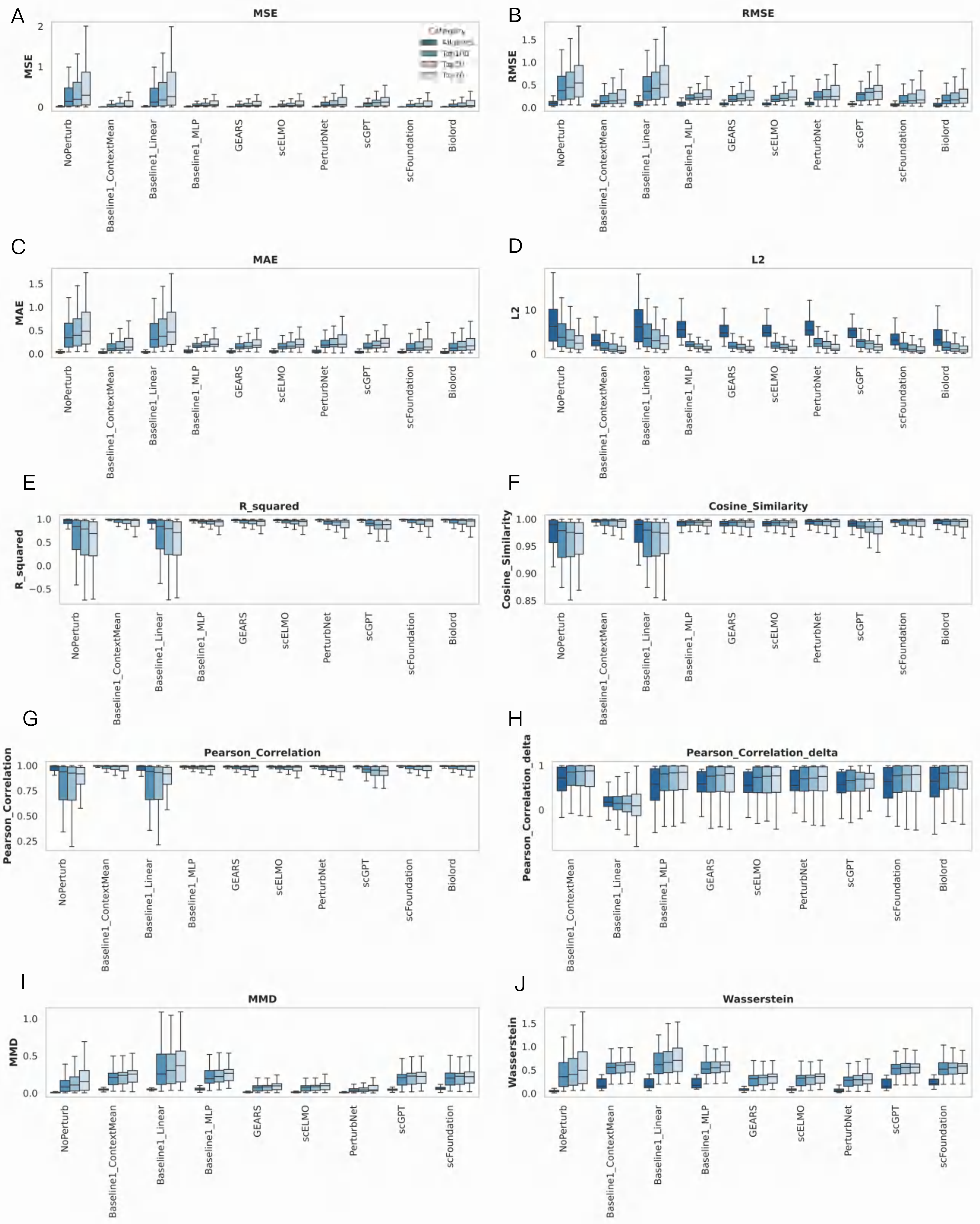

### fig. S3

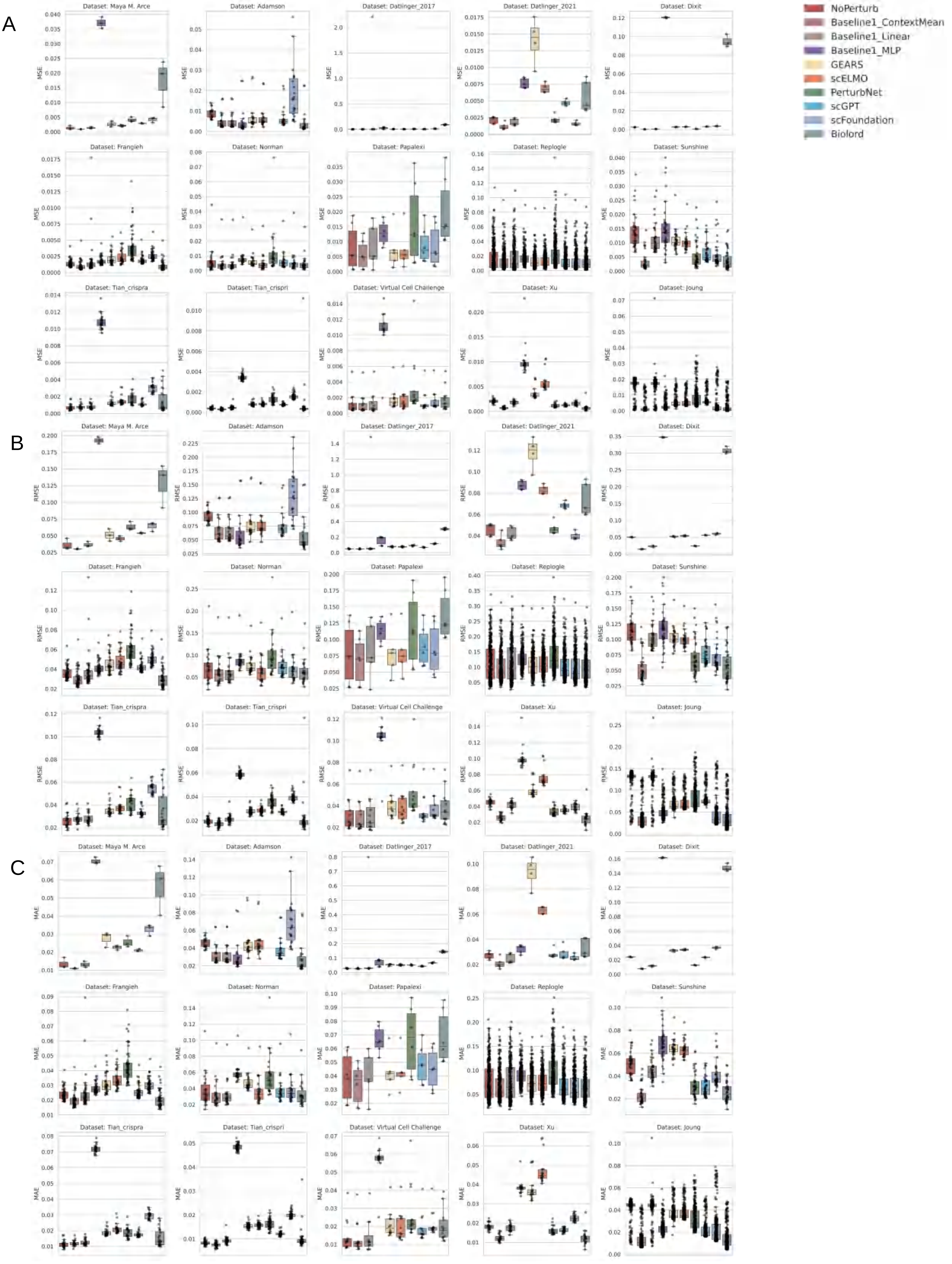

### fig. S4

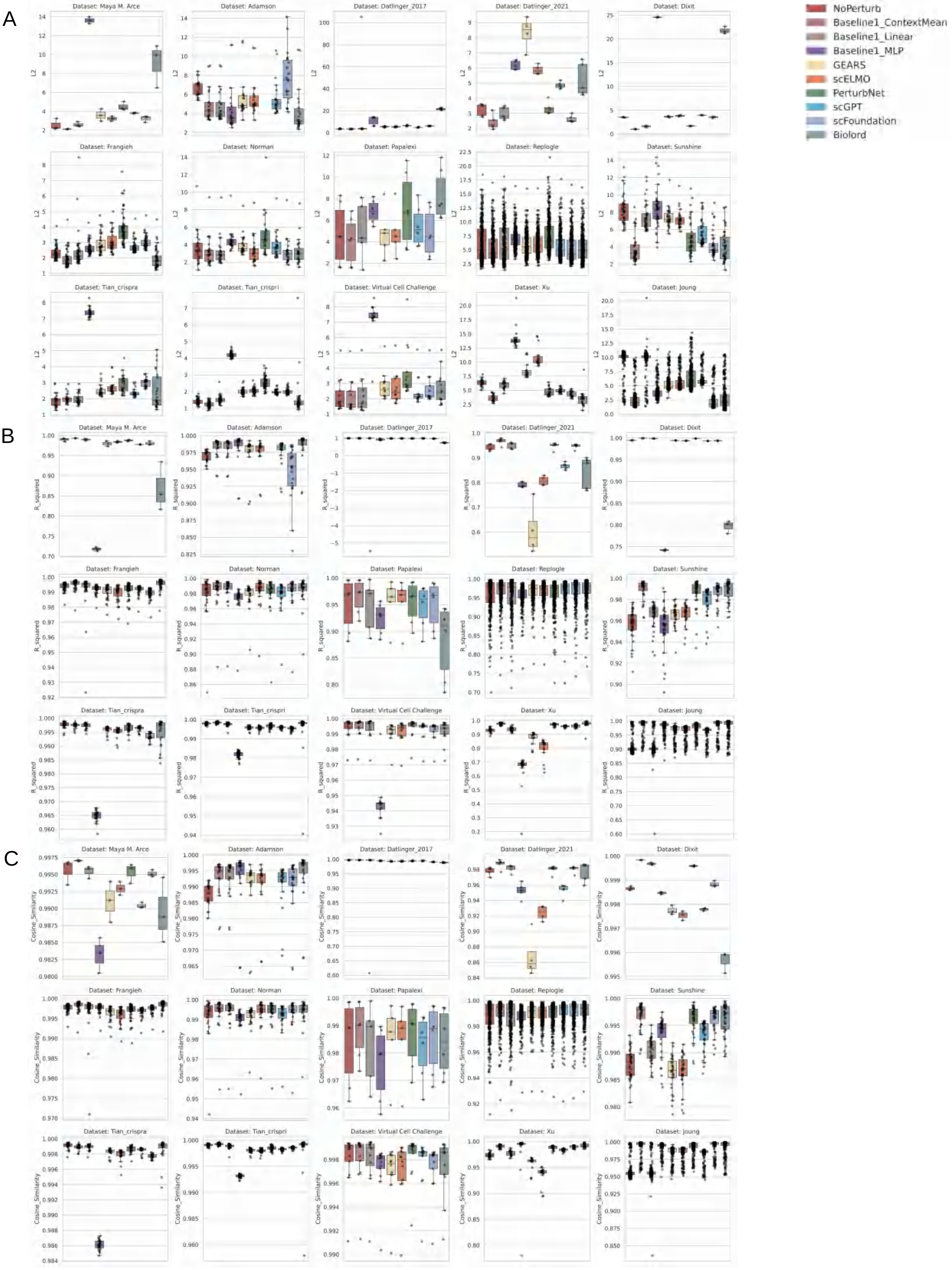

### fig. S5

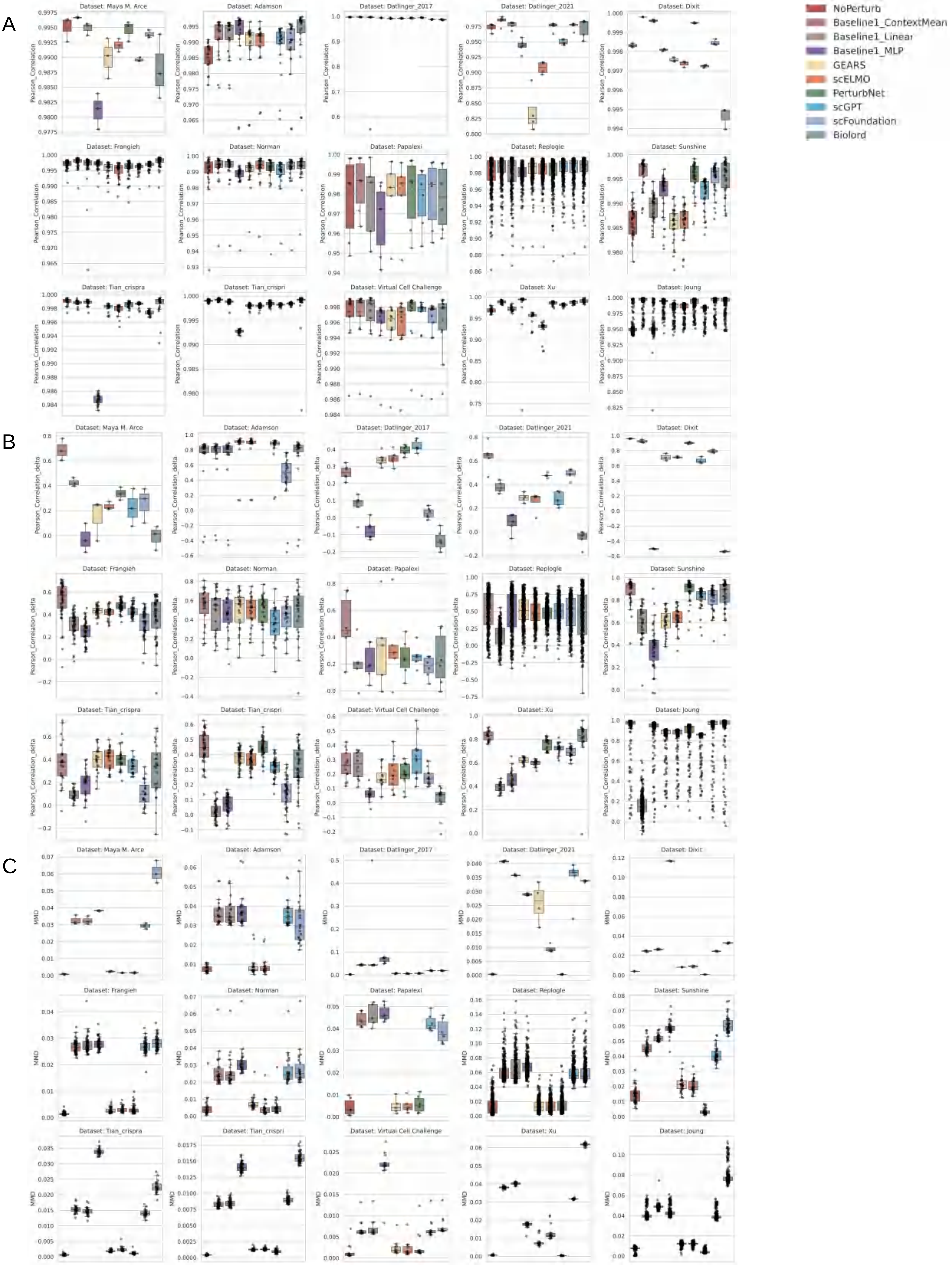

### fig. S6

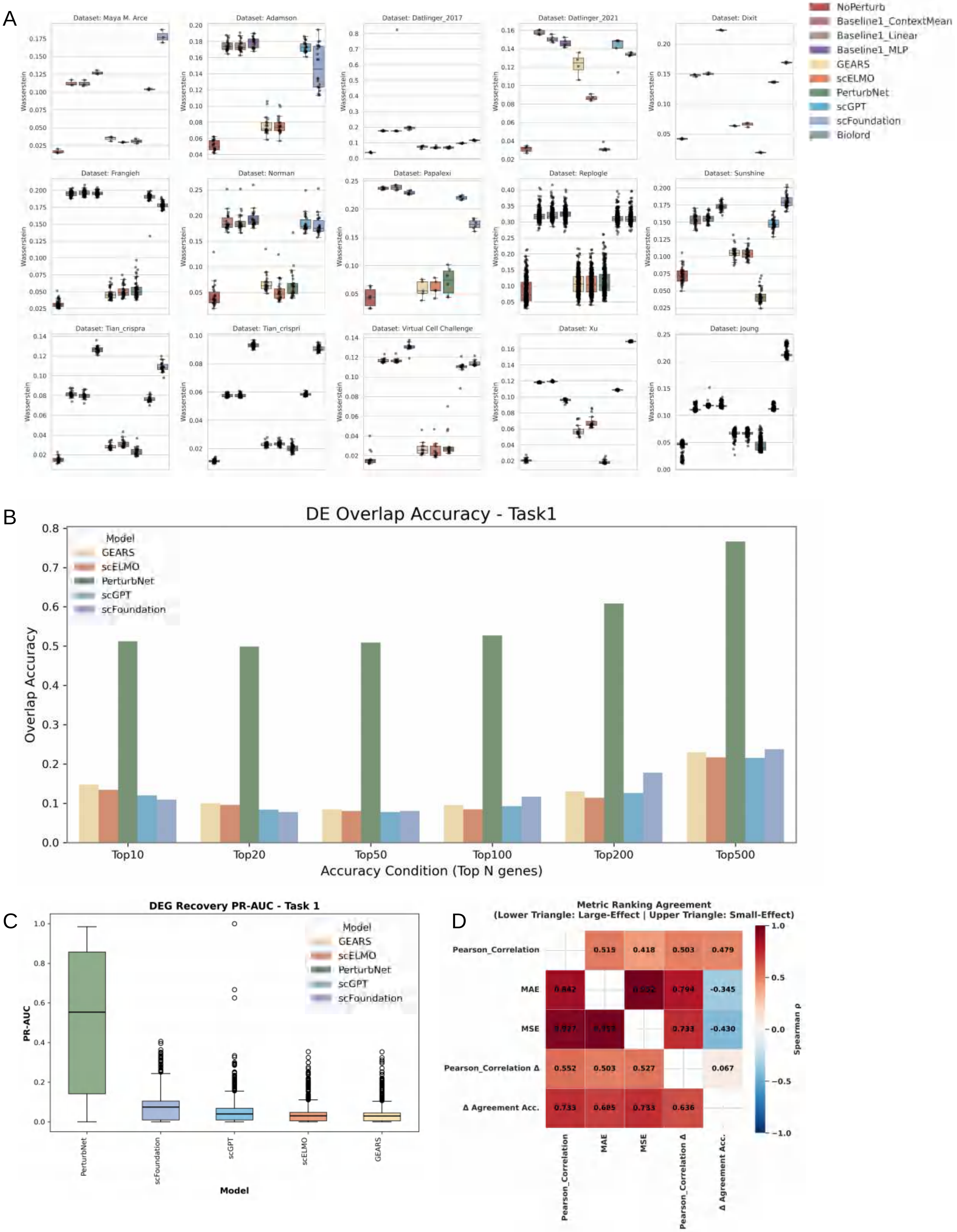

### fig. S7

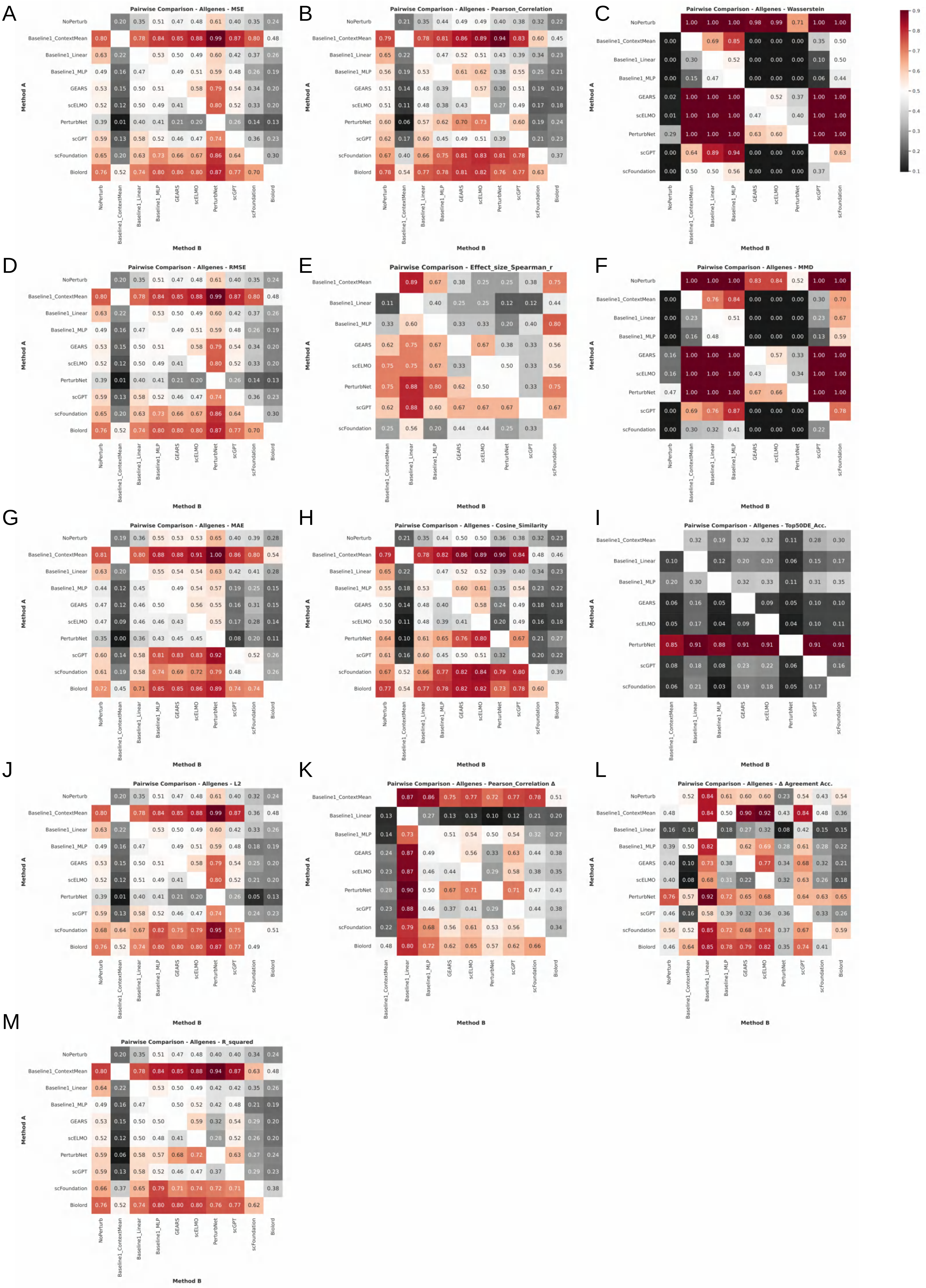

### fig. S8

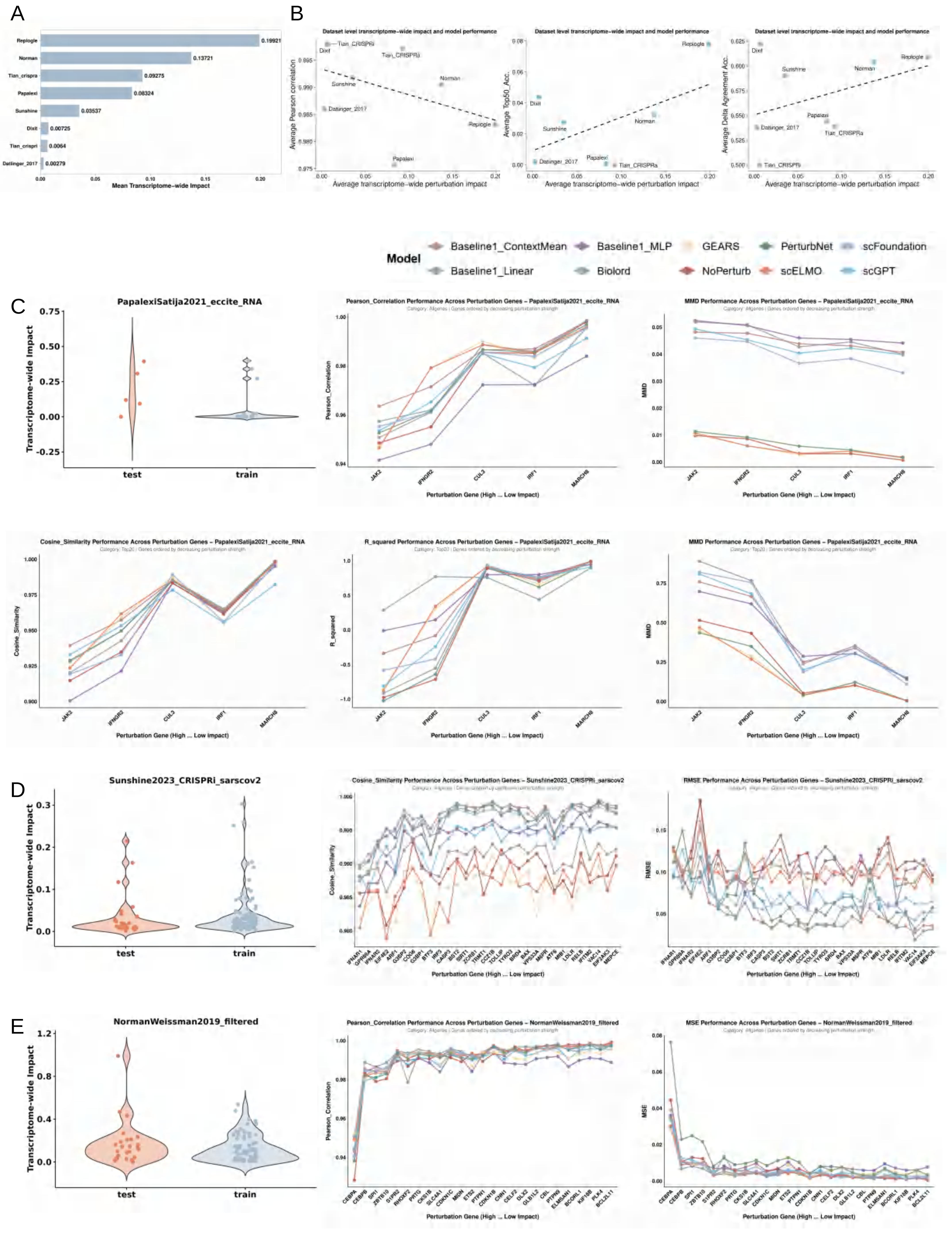

### fig. S9

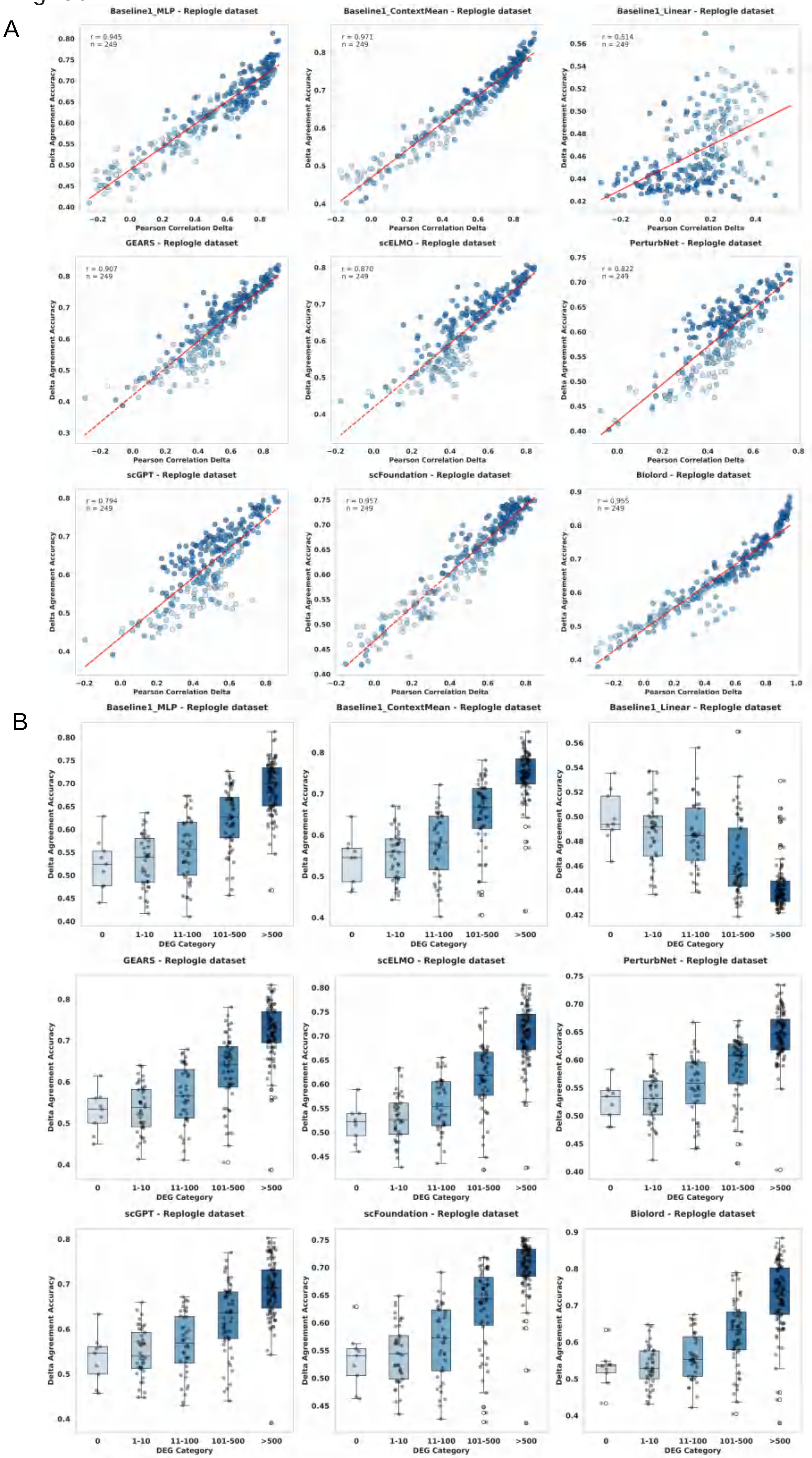

### fig. S10

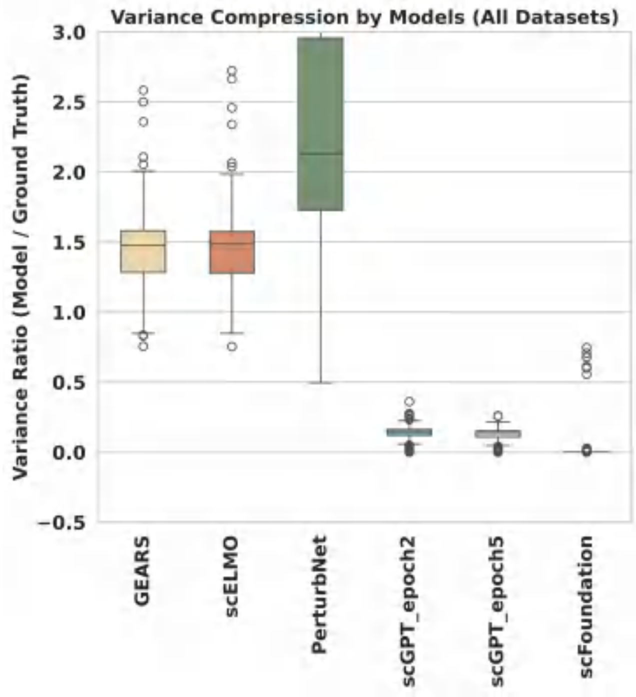

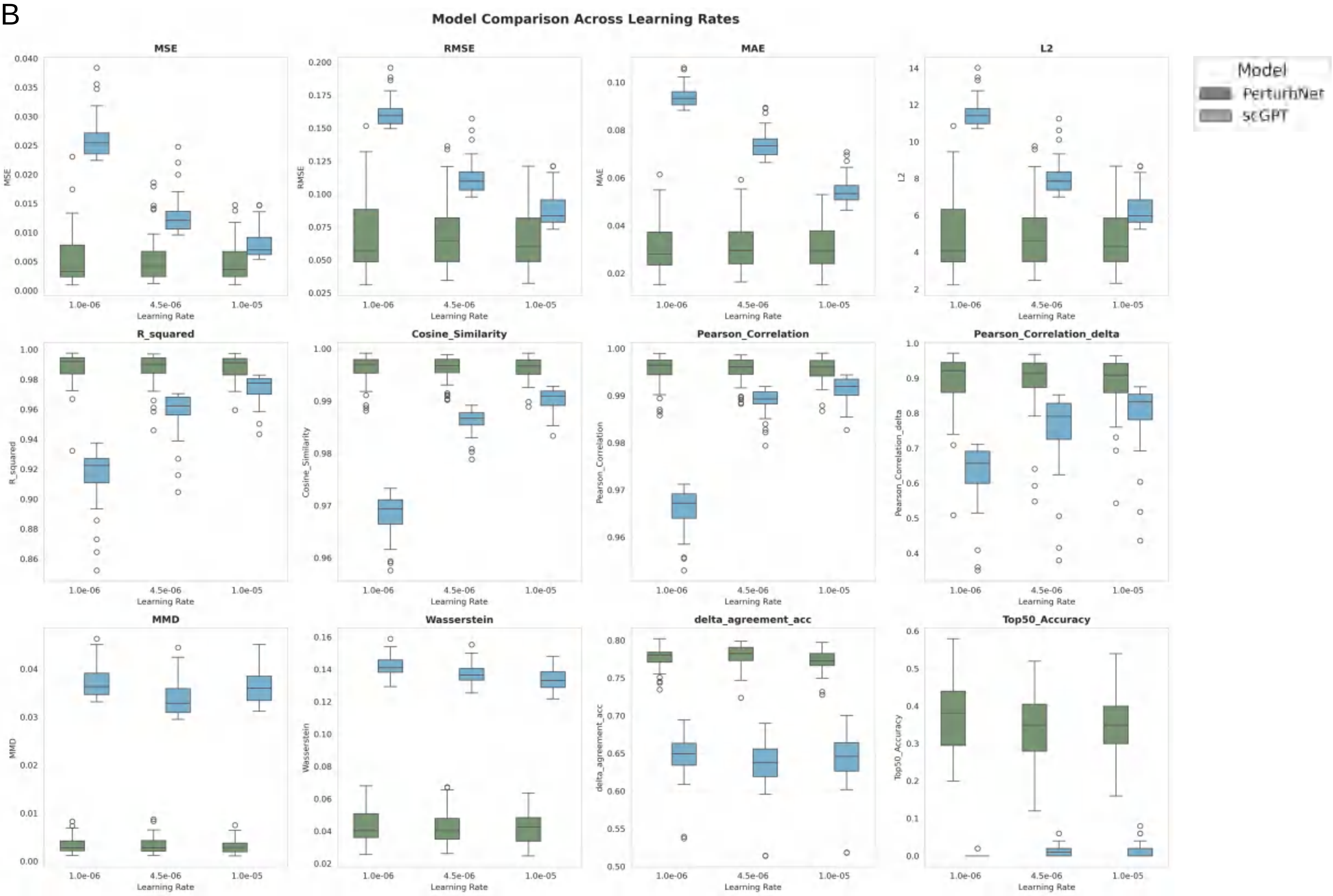

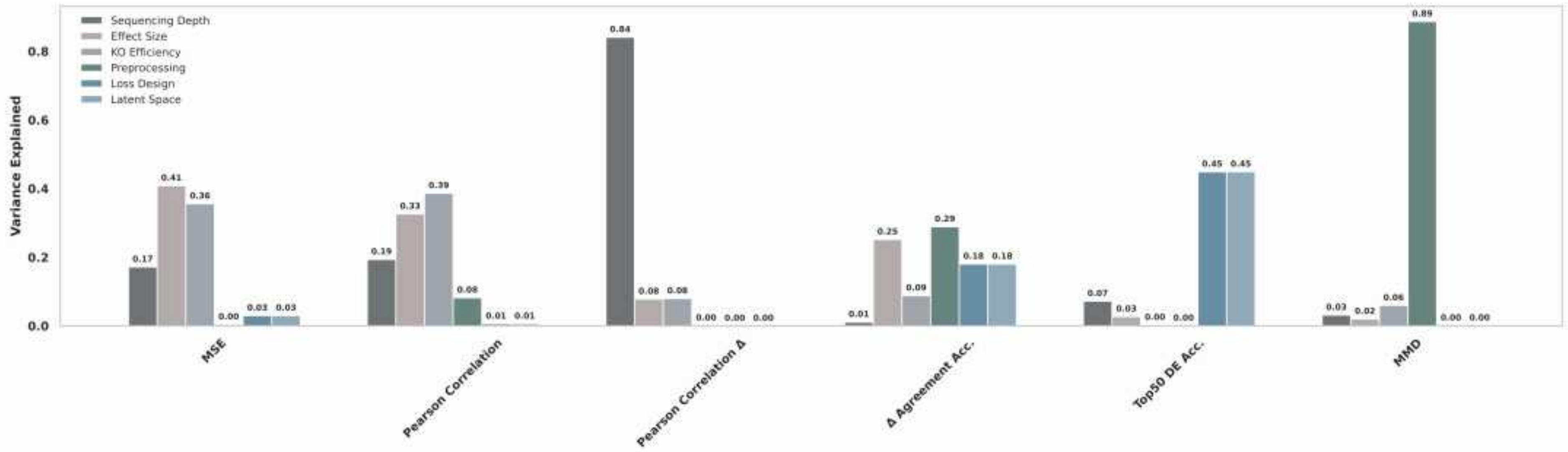

### fig. S11

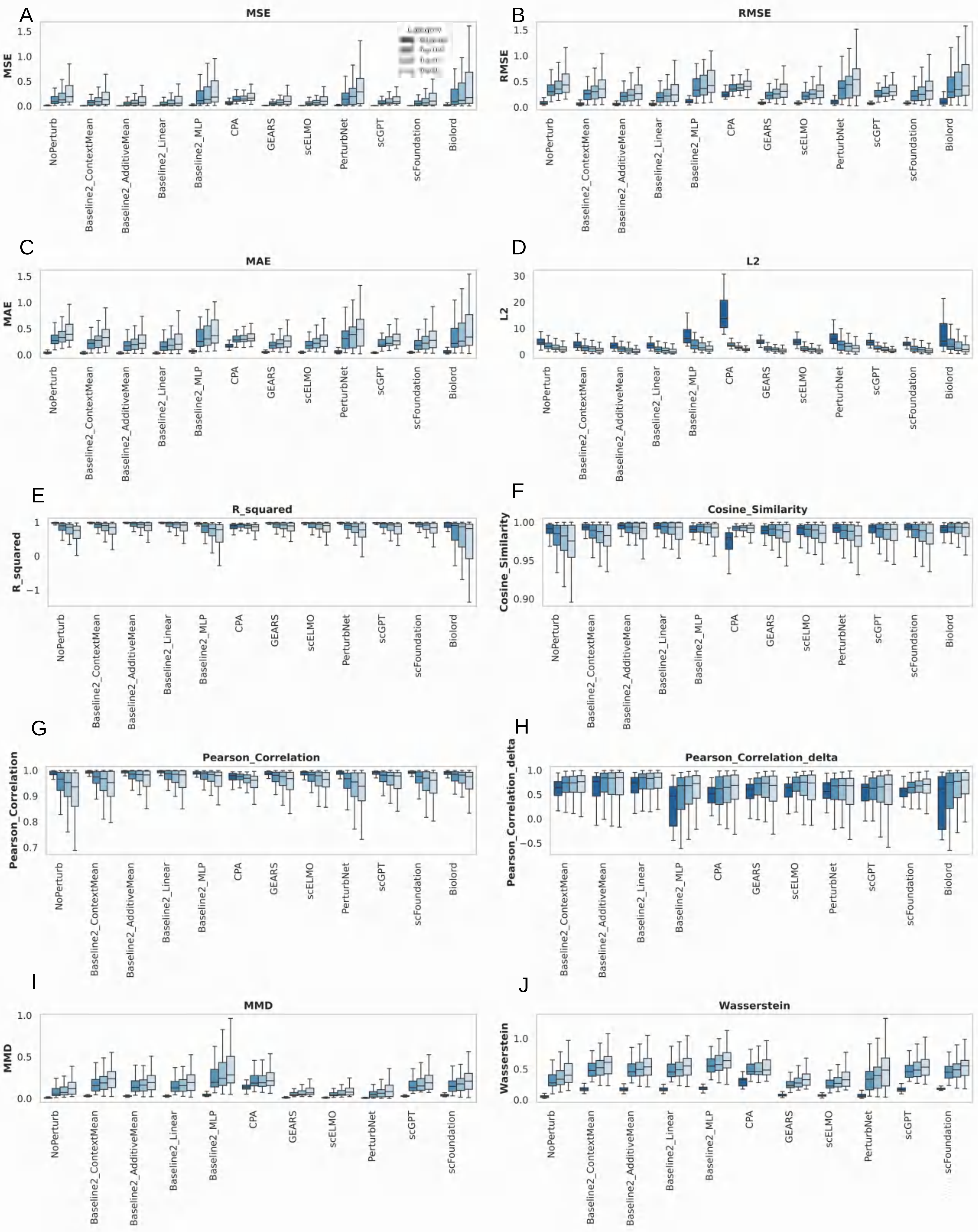

### fig. S12

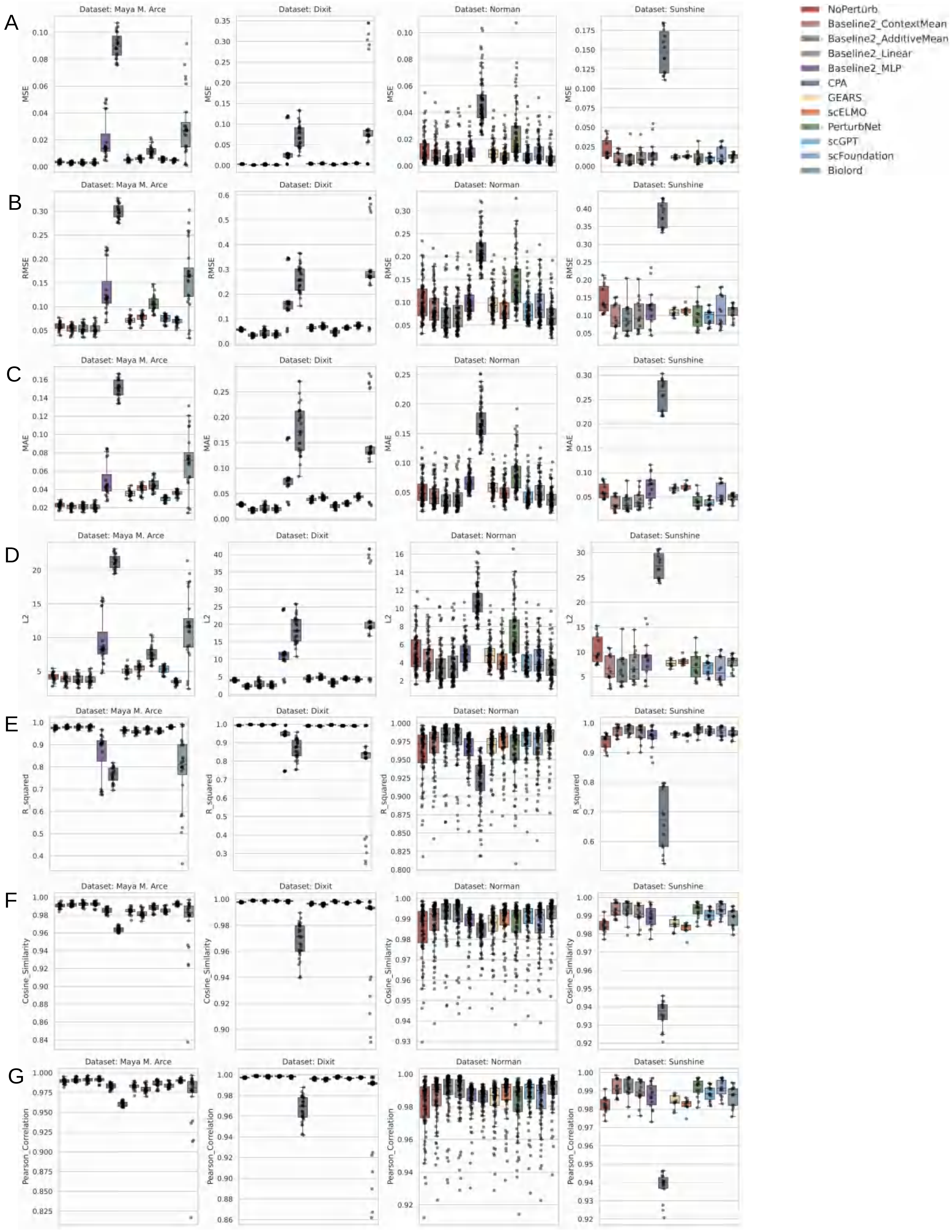

### fig. S13

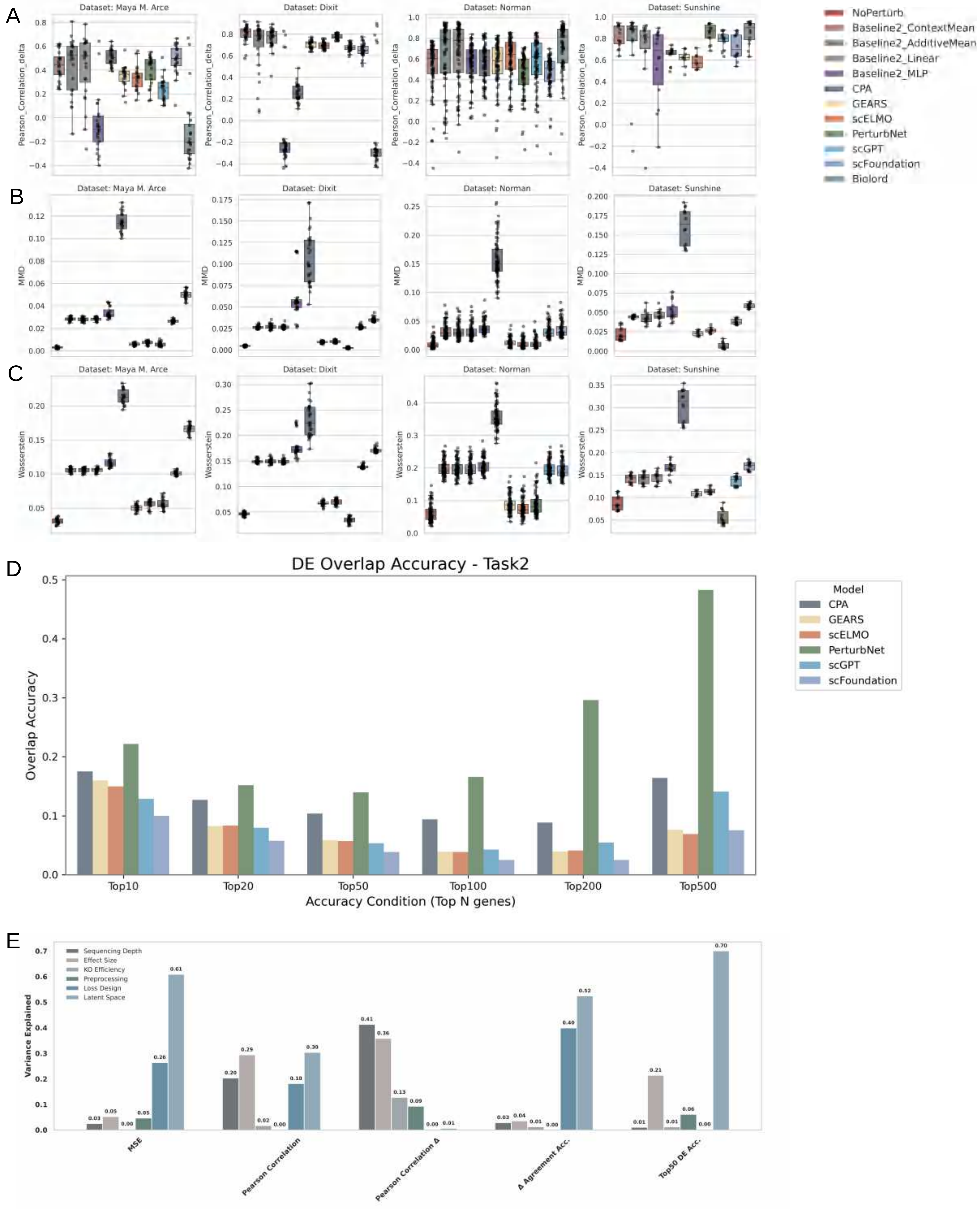

### fig. S14

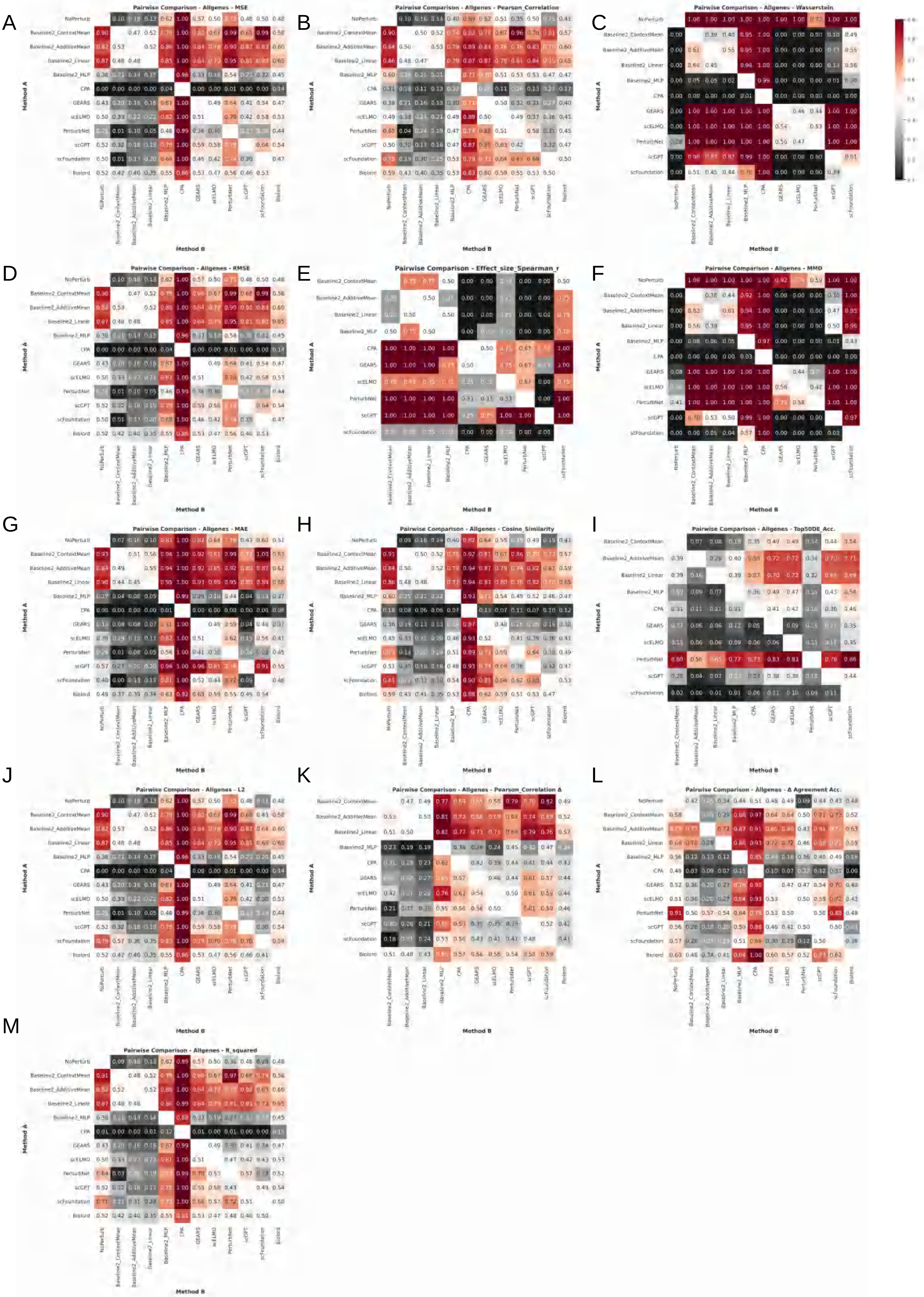

### fig. S15

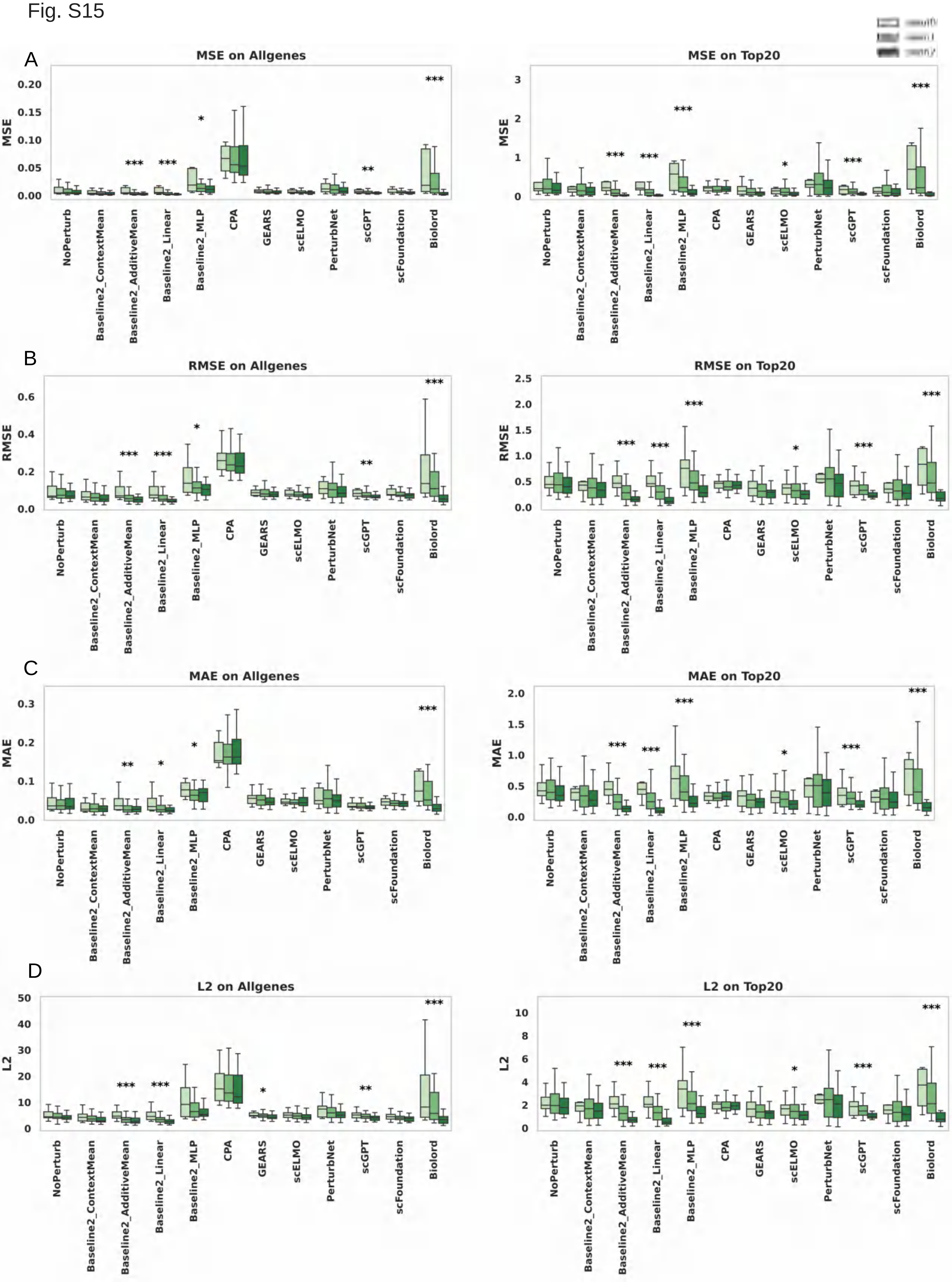

### fig. S16

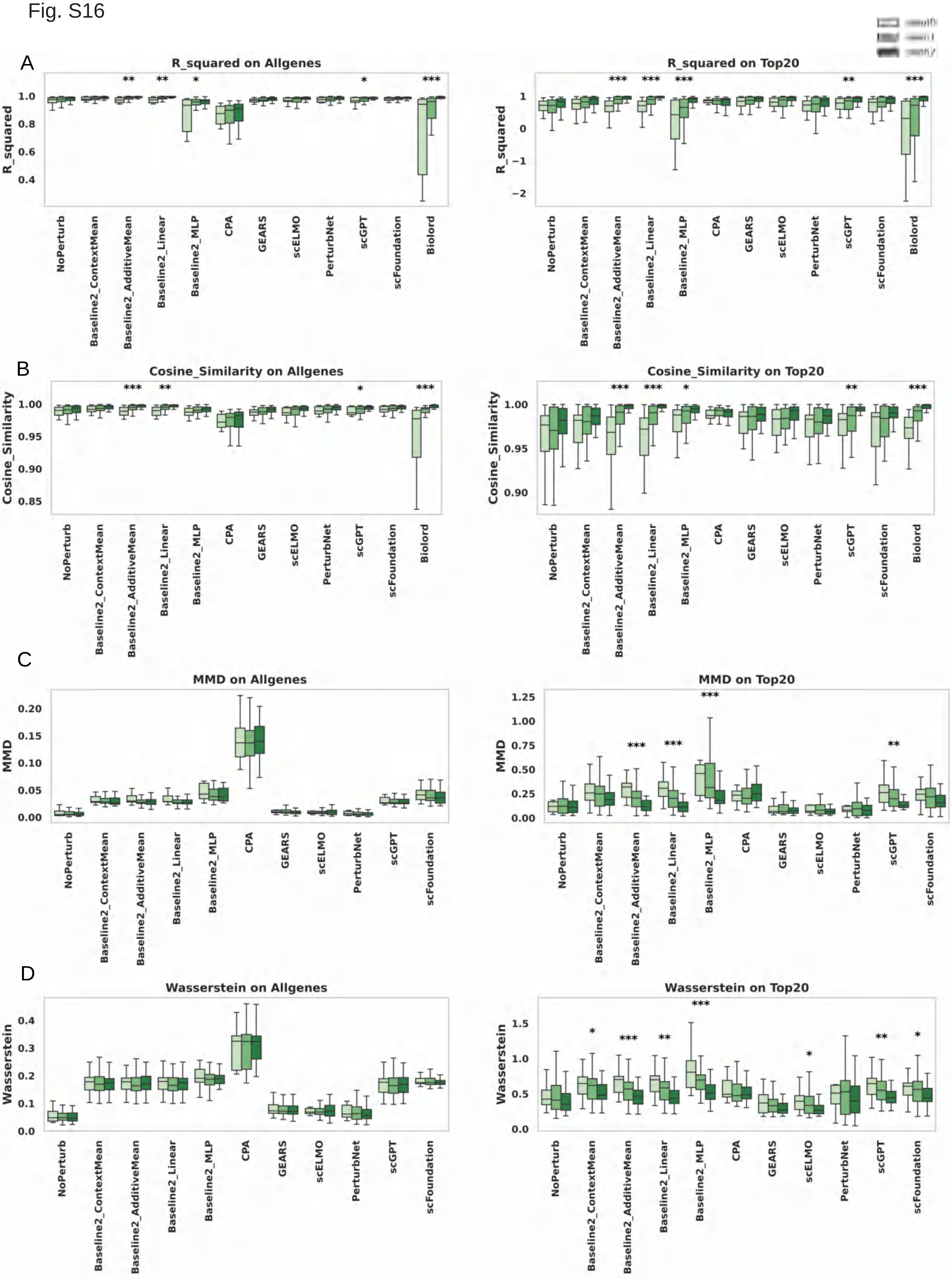

### fig. S17

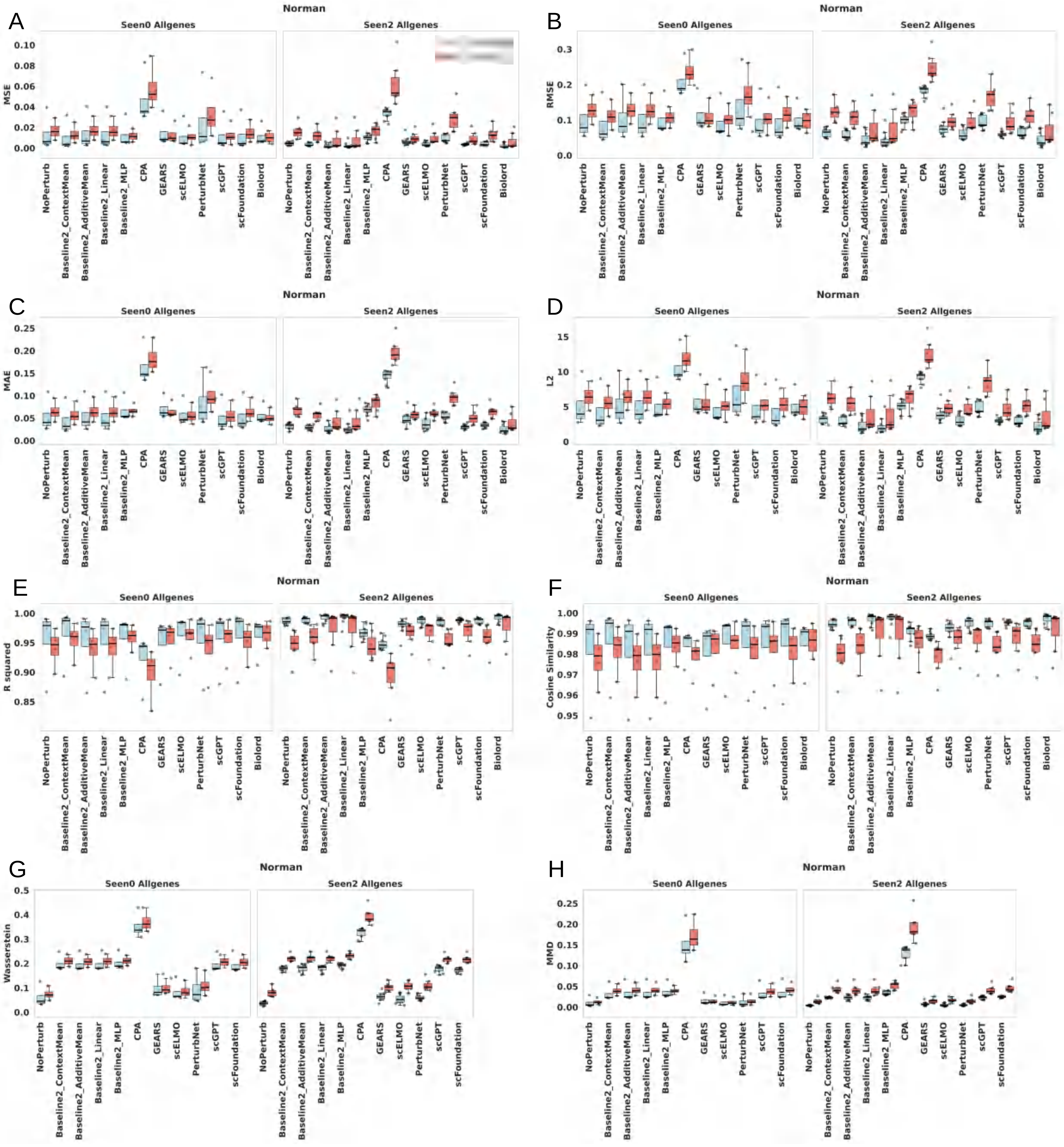

### fig. S18

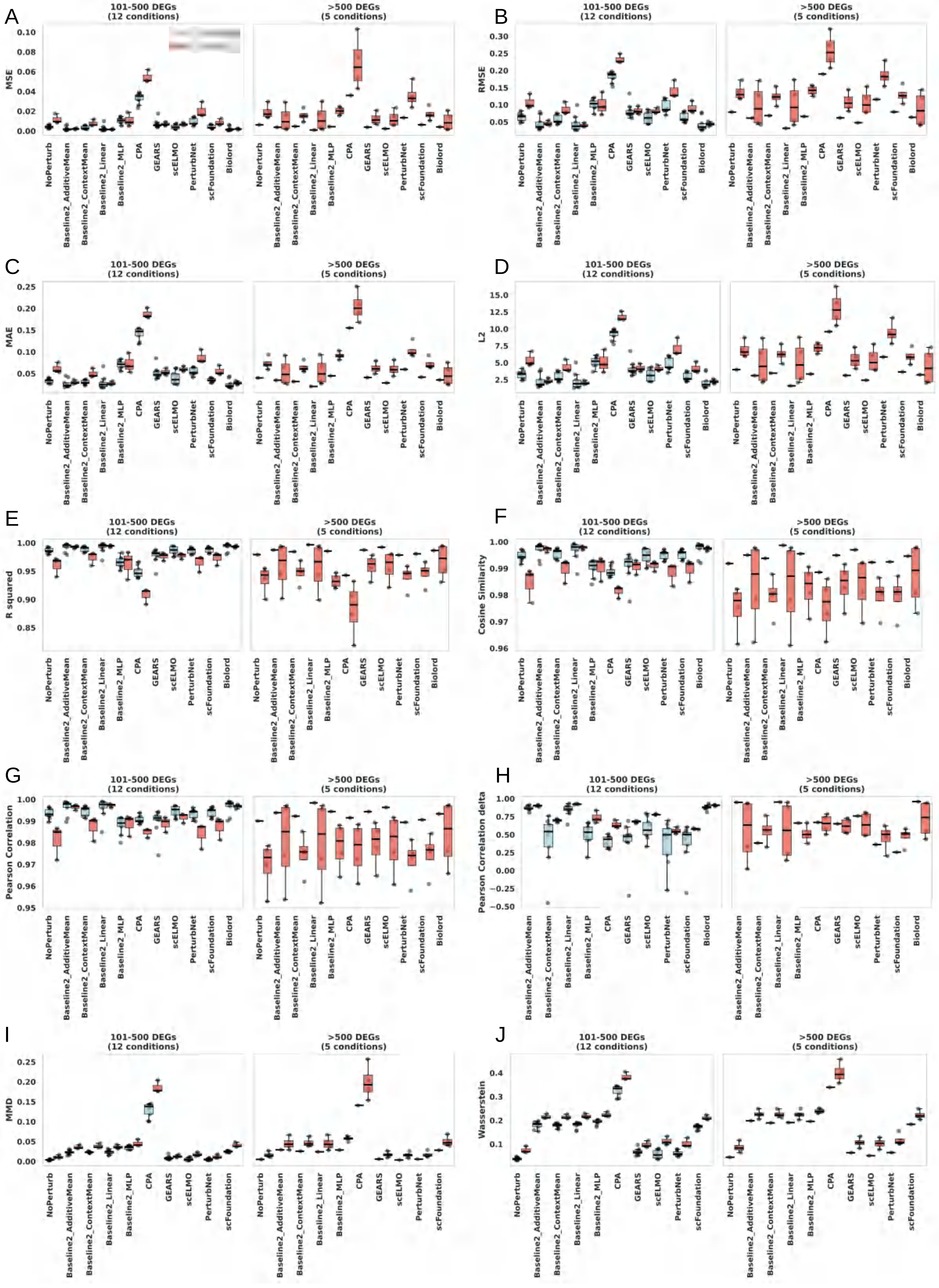

### fig. S19

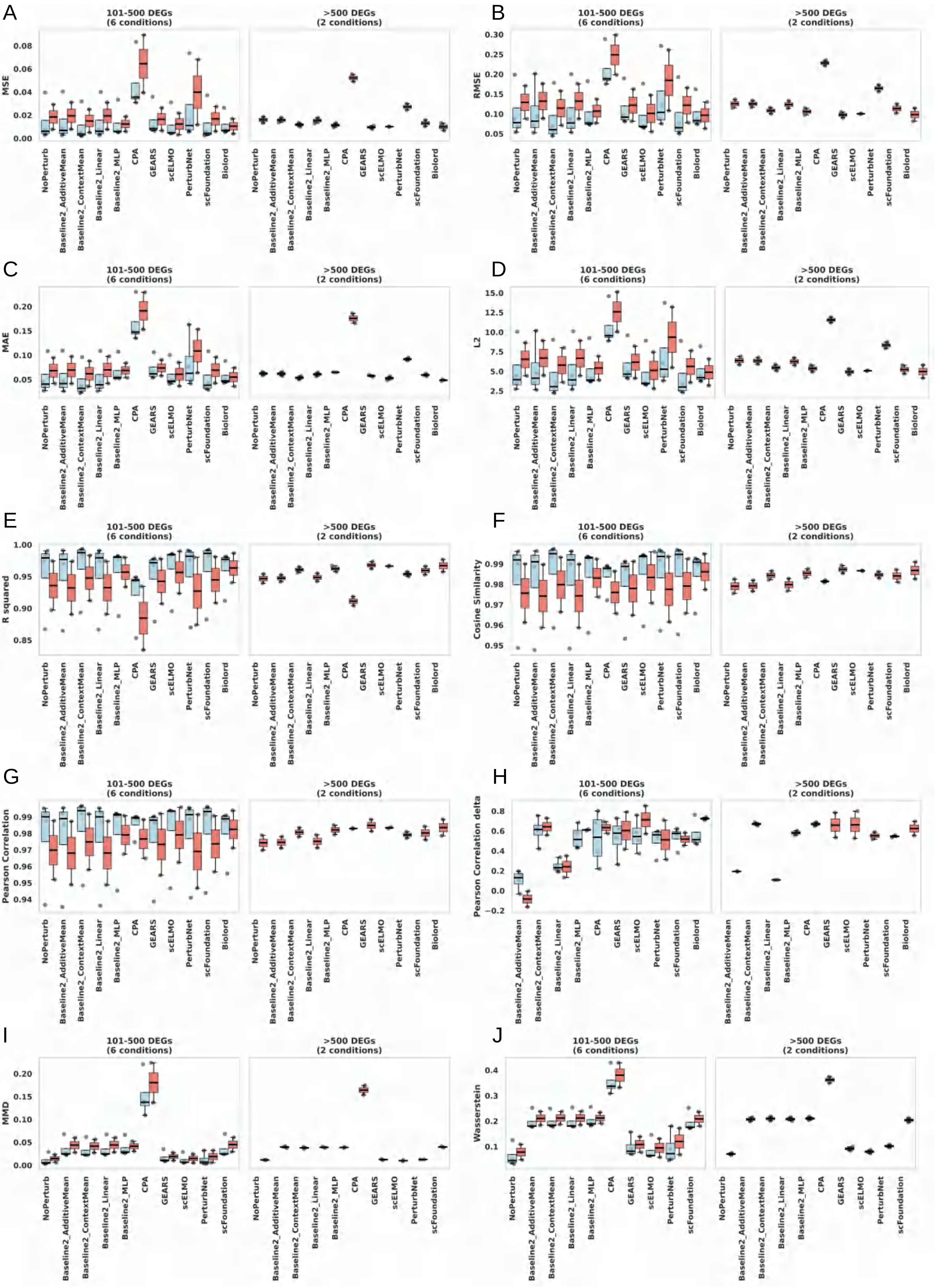

### fig. S20

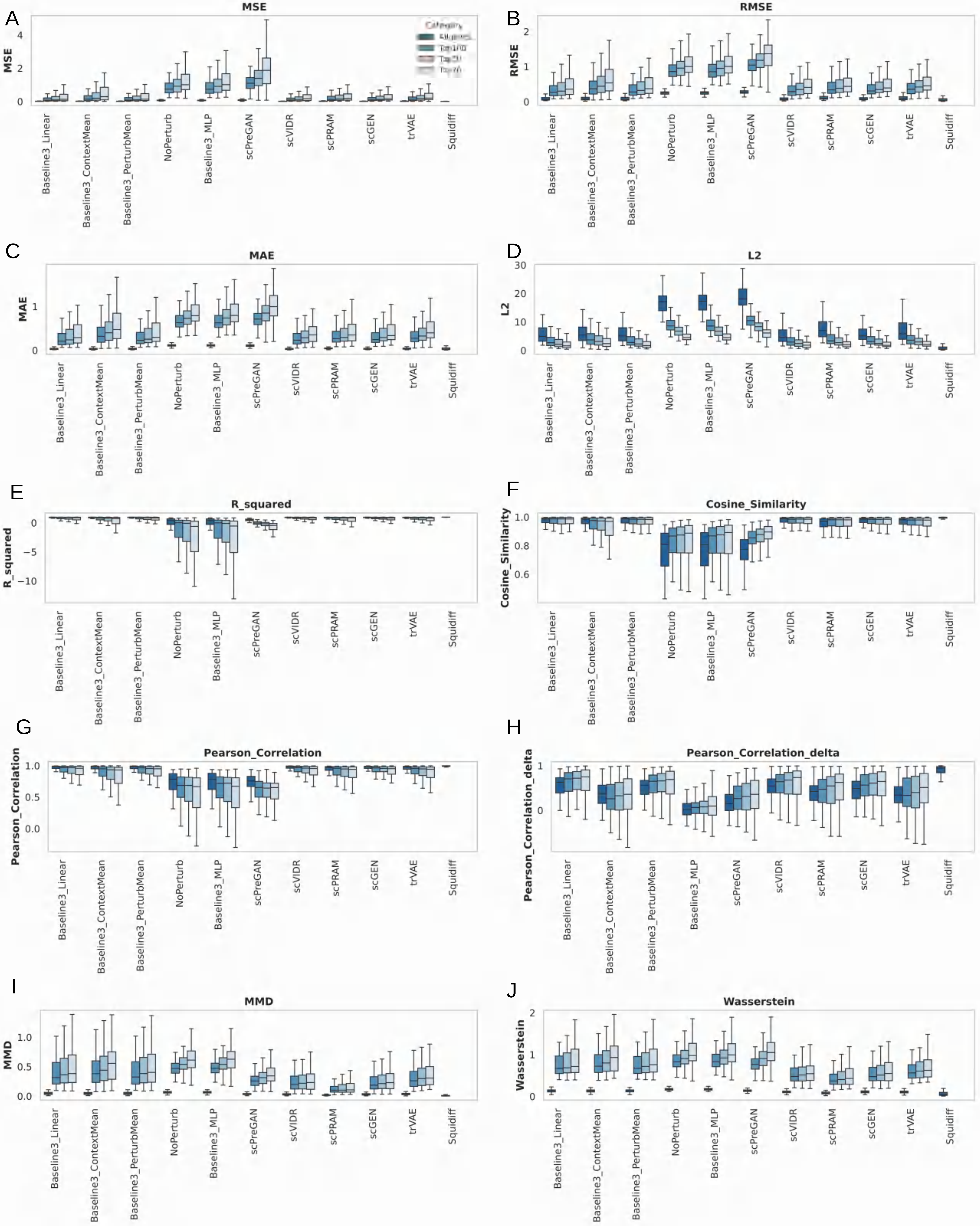

### fig. S21

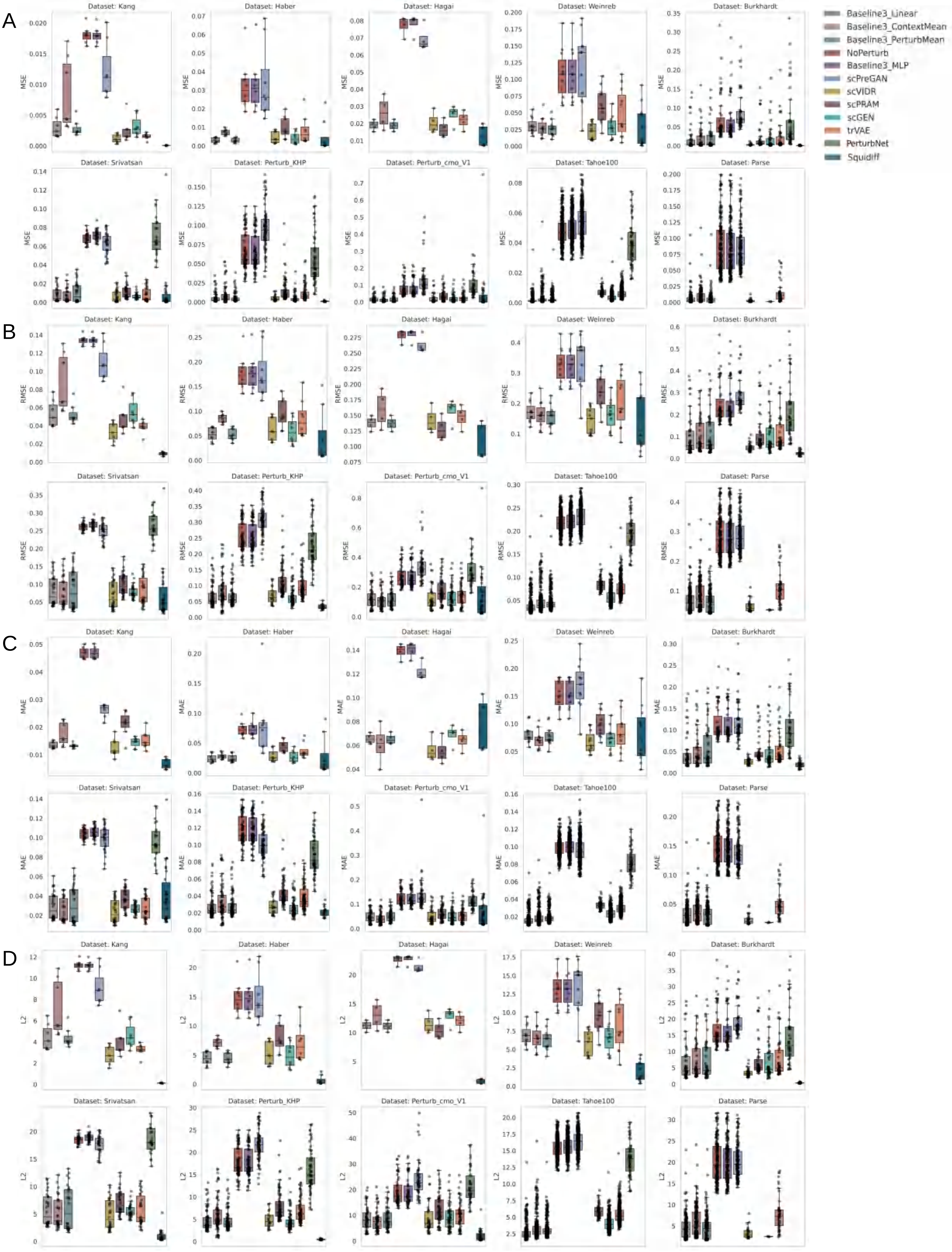

### fig. S22

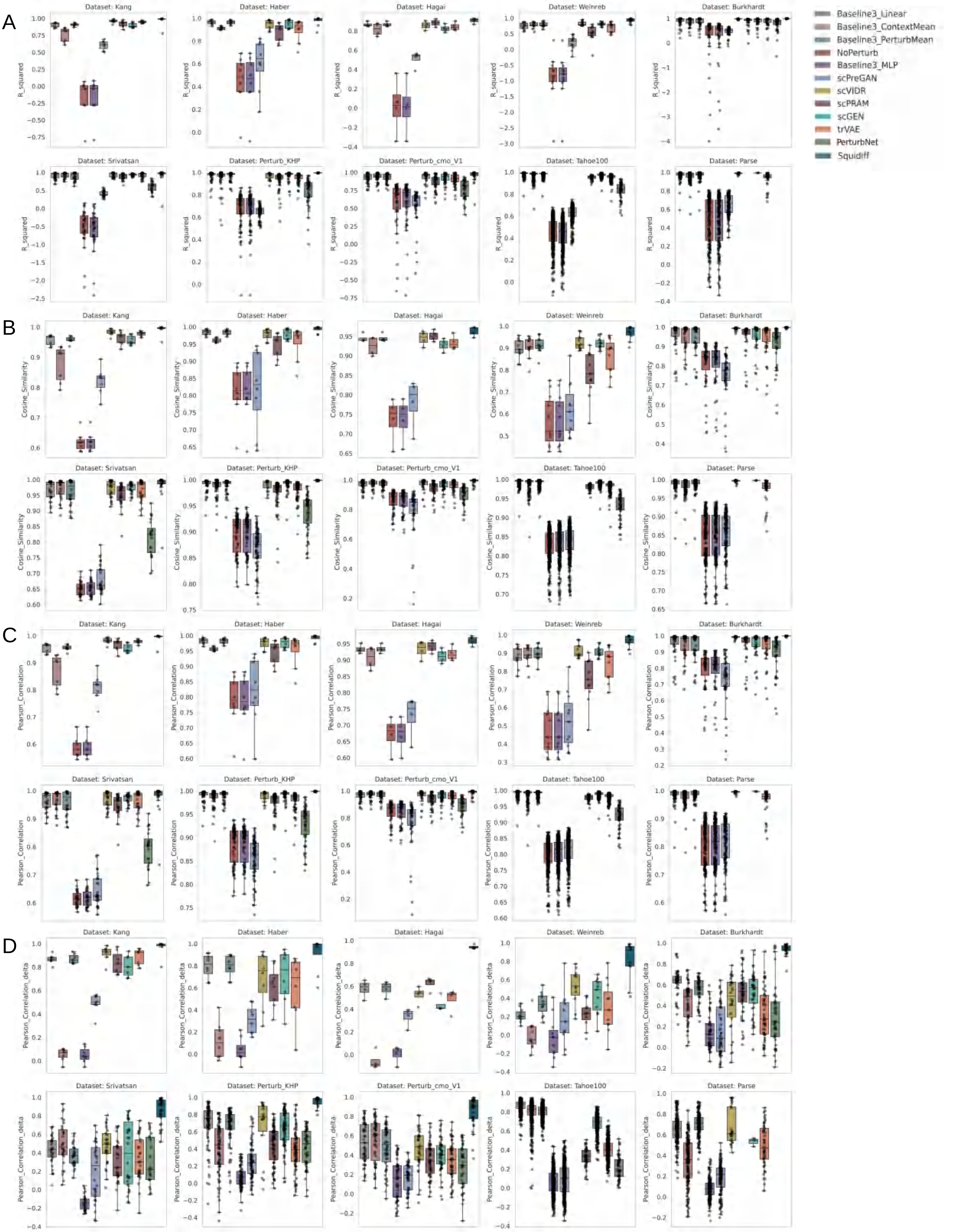

### fig. S23

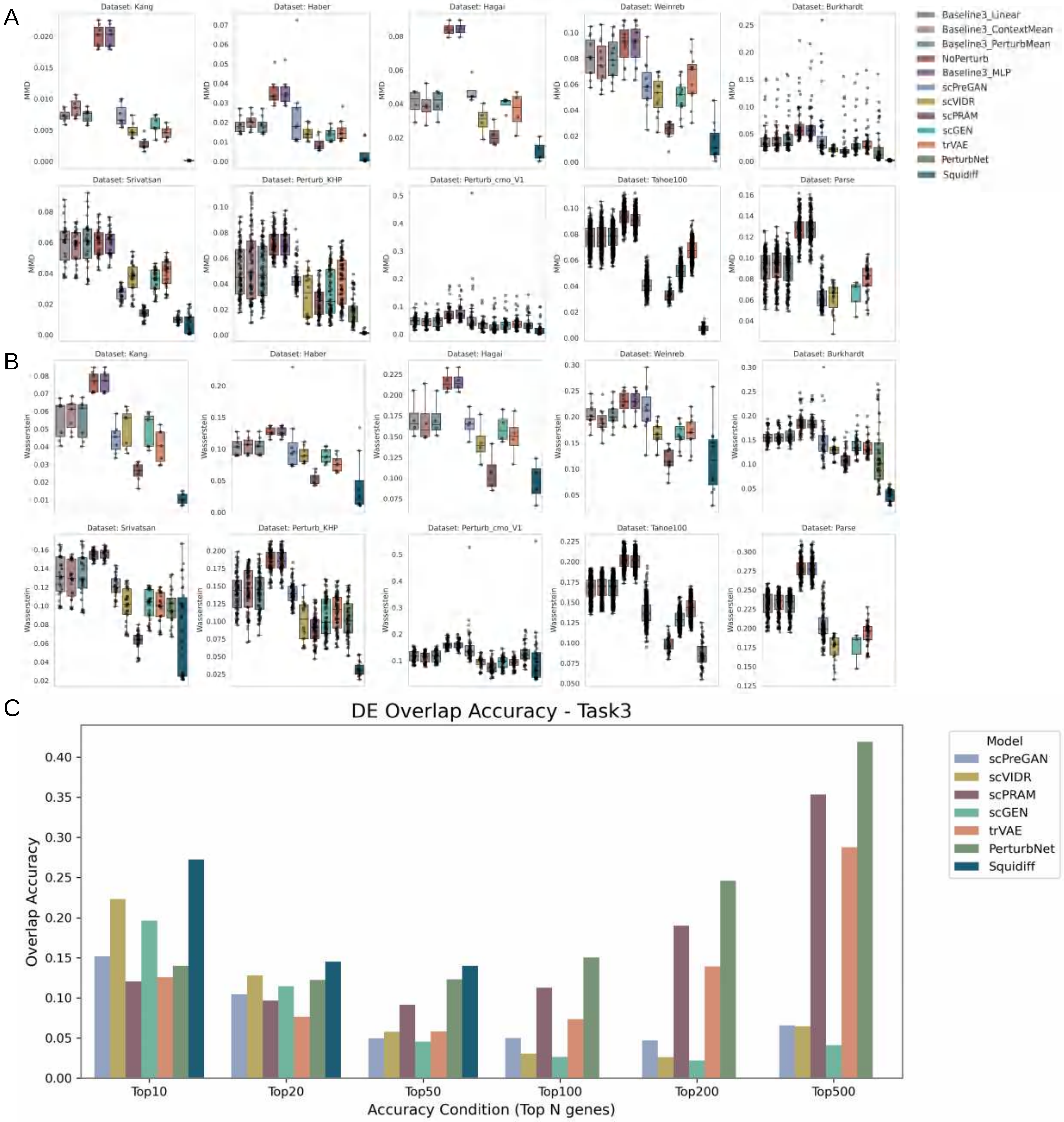

### fig. S24

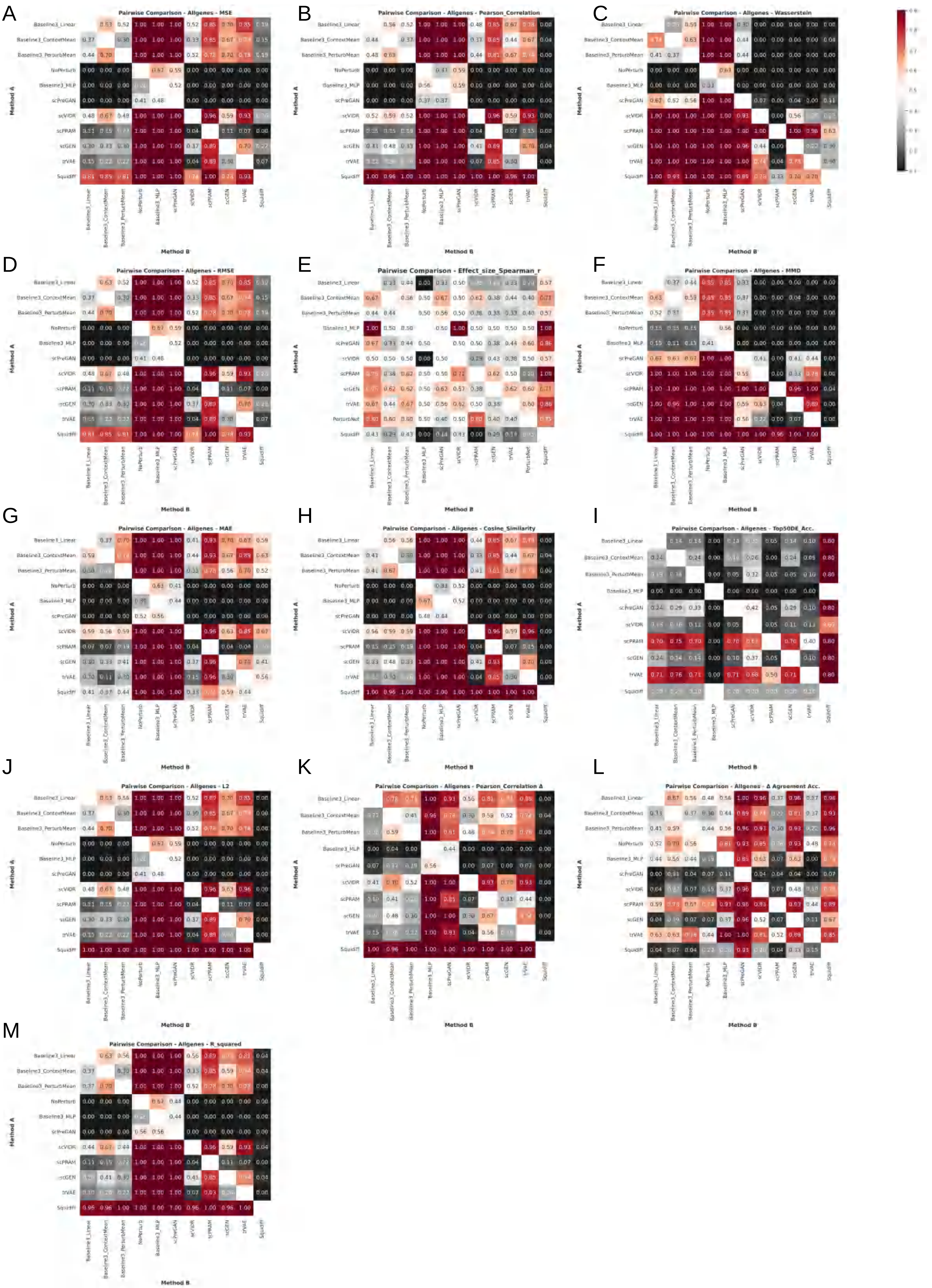

### fig. S25

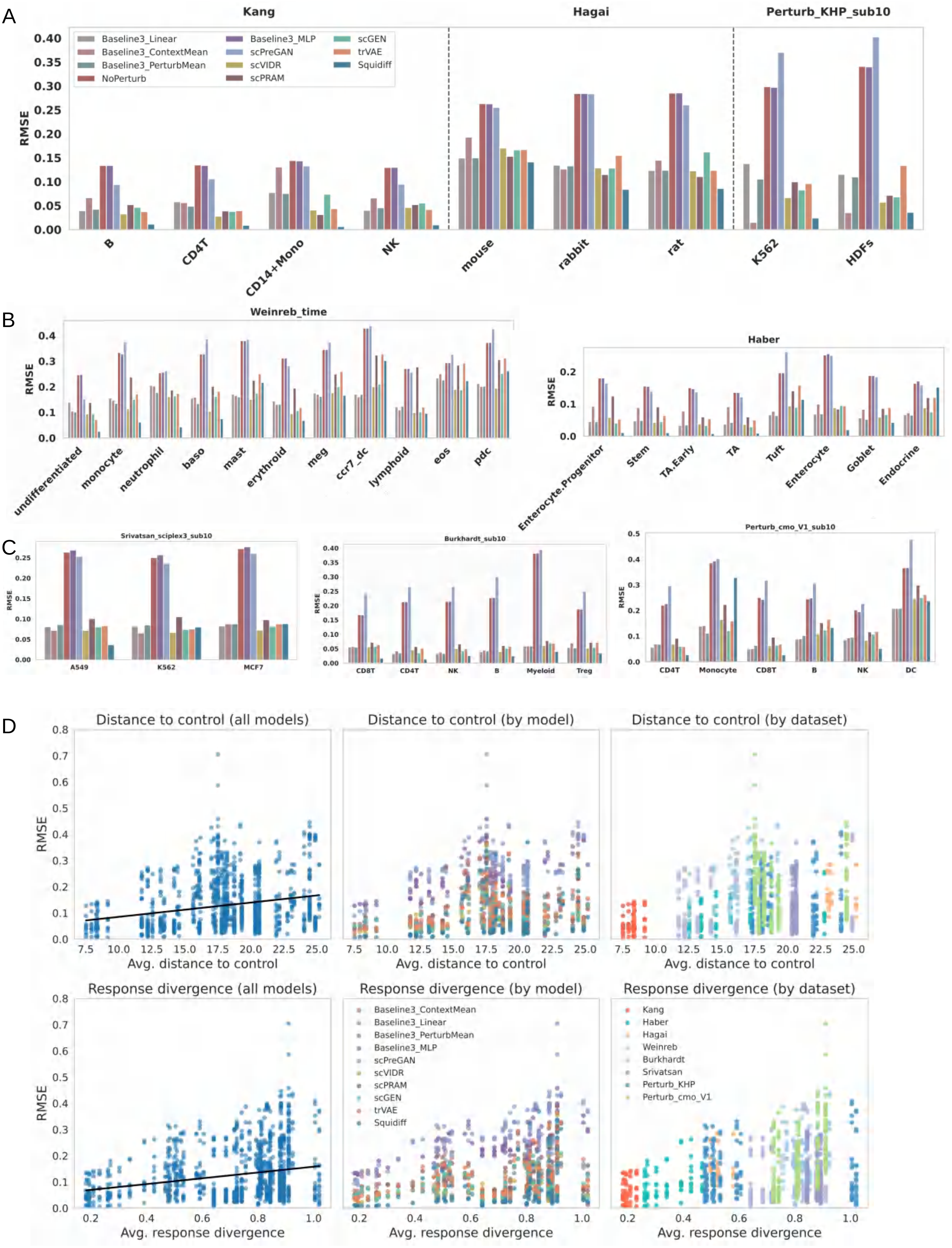

### fig. S26

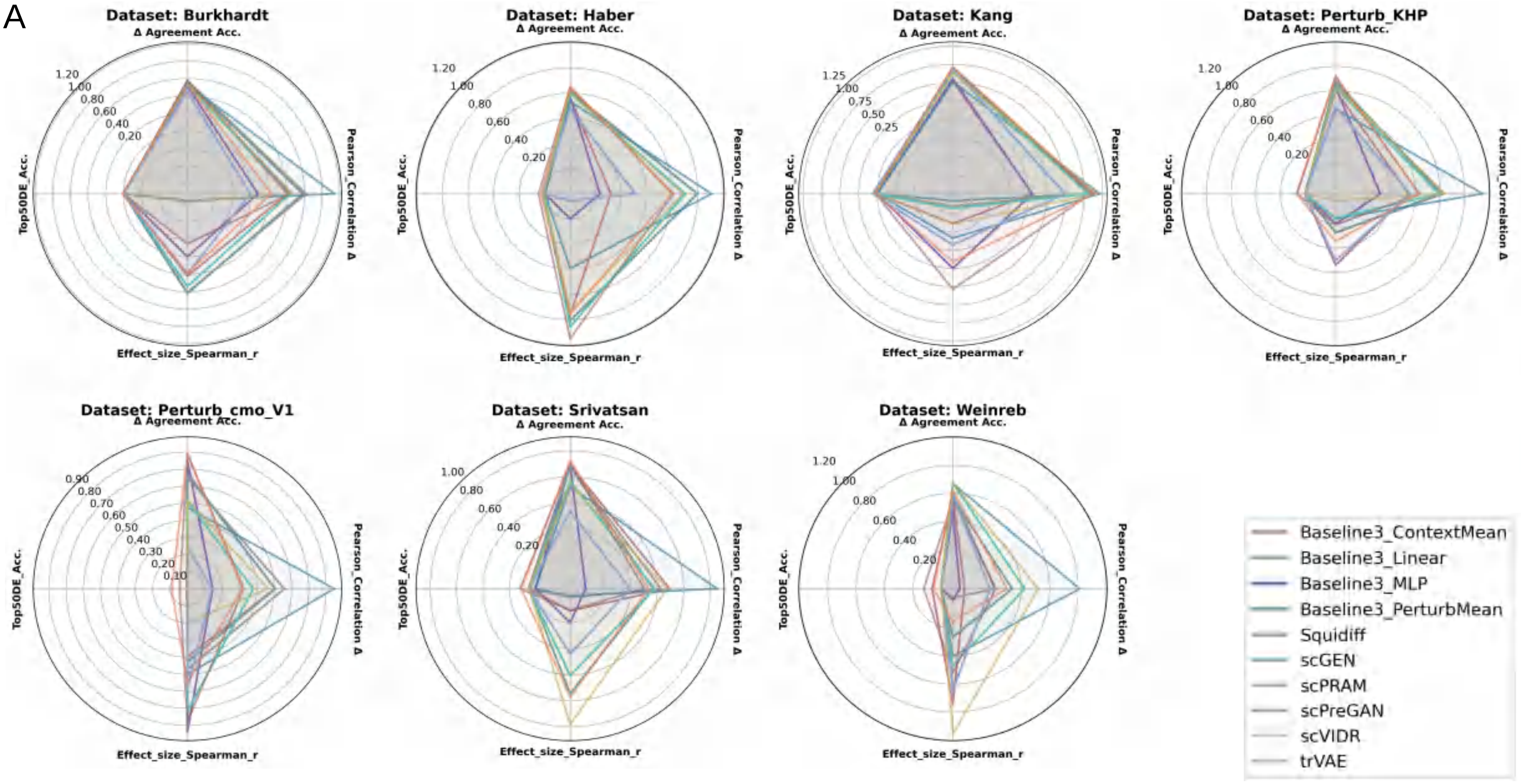

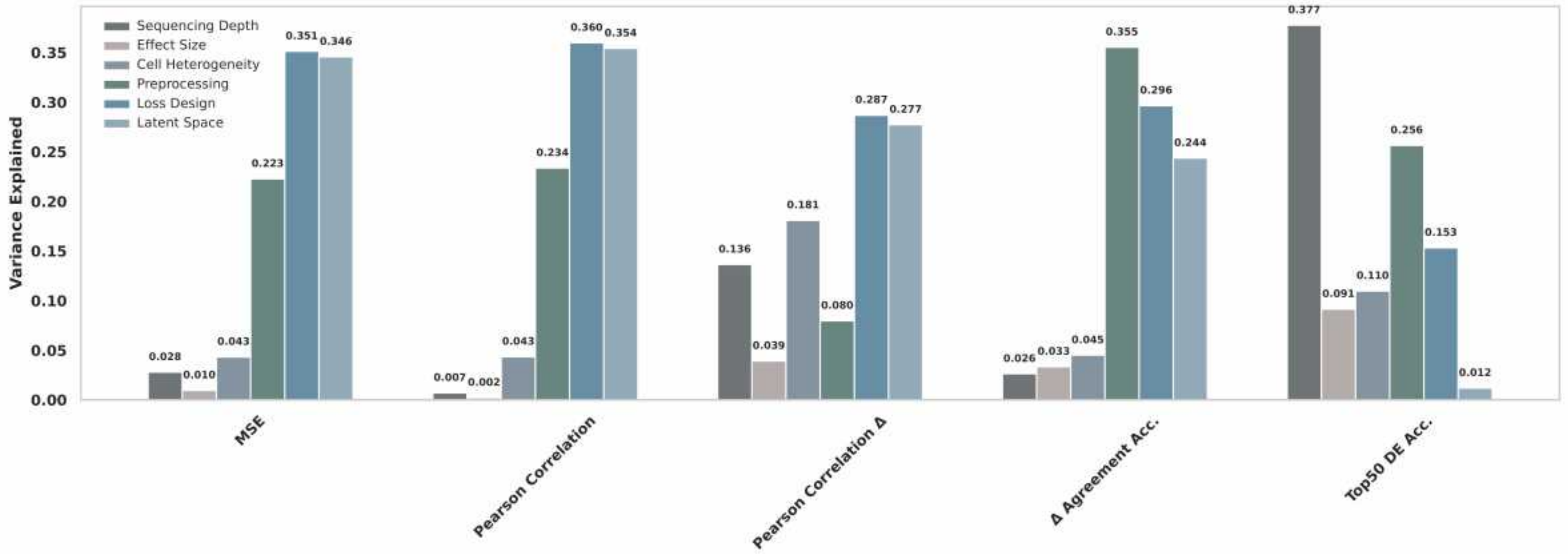

### fig. S27

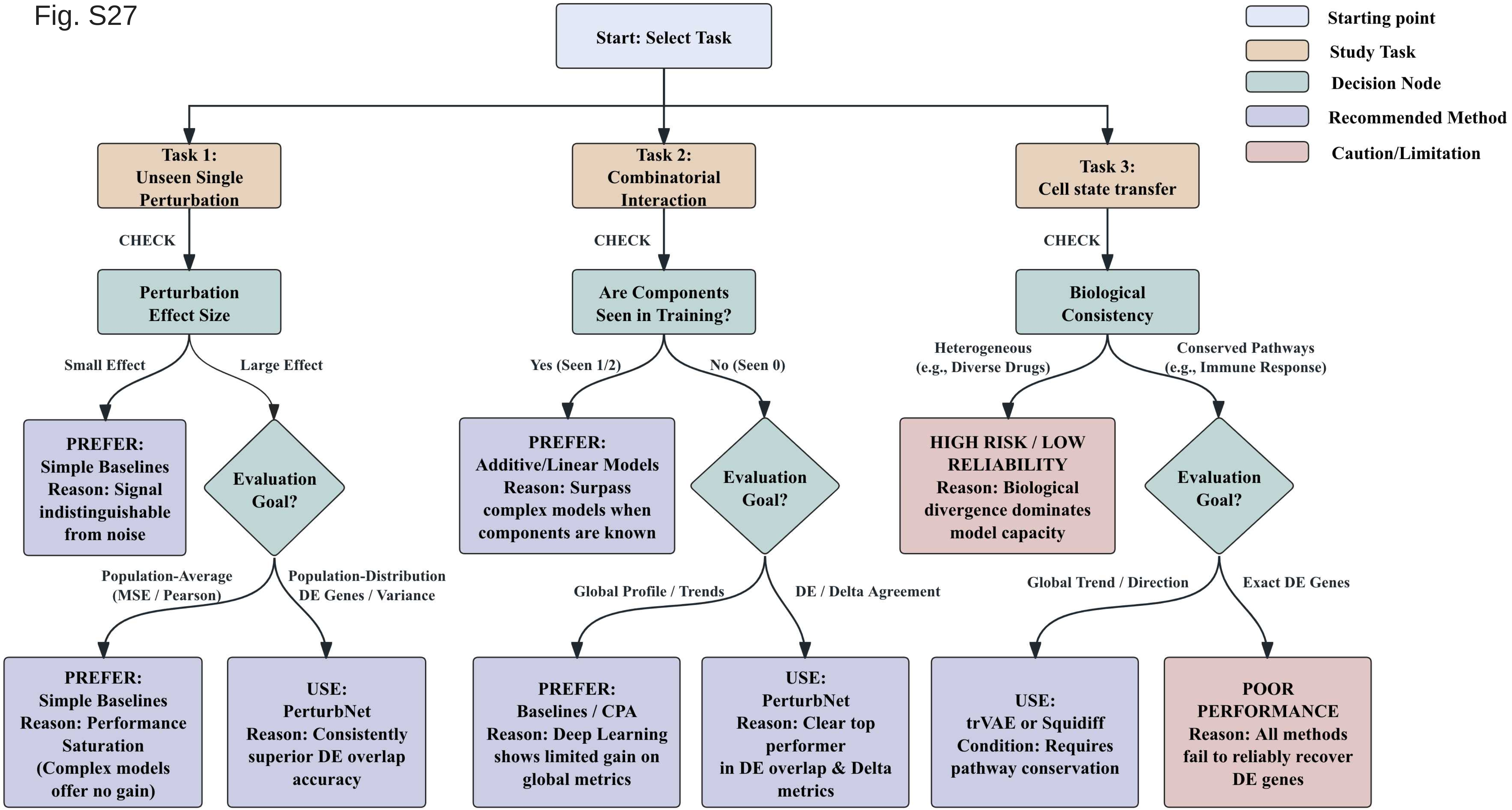
